# Integrative AI-Enabled Virtual Cell Modeling Reveals a Clinically Relevant Latent Effector State of Human CD8⁺ T Cells Undetectable by Conventional Analyses

**DOI:** 10.64898/2026.07.22.739582

**Authors:** Ying Li, Mojun Zhu, Roxana S. Dronca, Yiyi Yan, Wenjing Zhang, Yi Lin, Aaron S. Mansfield, Svetomir N. Markovic, Sean S. Park, Aubrey Y. Liew, Haidong Dong

## Abstract

Understanding how immune checkpoint inhibitors (ICIs) reshape human T-cell responses requires models that move beyond static transcriptomic snapshots and discrete cell-state classifications. Here, we present an integrative AI-enabled virtual cell framework that represents human CD8⁺ T-cell responses as dynamic and computable systems during ICI therapy. By integrating single-cell RNA sequencing with paired T-cell receptor sequencing within the C2S-scale foundation model, we construct a virtual representation of individual T cells, in which each cell is encoded by a unique functional identity that captures its transcriptional, signaling, and clonal characteristics. Using this framework, we identify a previously unrecognized dynamic latent effector state of CD8⁺ T cells characterized by intermediate expression of effector genes, distinct signaling activity, and ongoing clonal expansion. Across independent patient cohorts, the virtual cell model consistently indicates that ICI therapy mainly acts by unmasking pre-existing effector potential rather than inducing de novo effector differentiation. Notably, this latent effector population remains transcriptionally restrained despite active signaling and clonal expansion, revealing a hidden reservoir of antitumor immune capacity. More broadly, our study demonstrates how AI-enabled virtual cell modeling can reconstruct latent cellular states and their dynamic transitions from multidimensional single-cell data. By incorporating functional identity into virtual cell model, this framework uncovers biologically meaningful yet non-obvious T-cell effector program during cancer immunotherapy and provides a generalizable approach for studying immune dynamics in human disease.

## Introduction

Efforts to develop computational virtual cell models capable of predicting cellular behavior across diverse biological contexts have accelerated rapidly in recent years^1–3^. Advances in large-scale foundation models trained on massive single-cell datasets have enabled increasingly powerful representations of cellular states^4,5^. However, despite these advances, substantial conceptual and technical barriers remain. In particular, recent studies highlight that model scaling alone is insufficient to overcome a central limitation: the lack of context diversity in current training data^6^. Most existing models achieve strong performance within narrowly defined biological settings but fail to generalize across timepoints, treatments, and patient-specific conditions. This limitation reflects a deeper challenge of causal transportability, whereby models trained on restricted data distributions cannot reliably deduce to clinically relevant situations. Consequently, current virtual cell frameworks remain constrained in their ability to deliver mechanistically interpretable and clinically predictive insights, particularly in complex therapeutic settings.

To address these limitations, we sought to develop a context-aware virtual cell framework that integrates large-scale representation learning with mechanistic modeling. We selected the C2S-scale foundation model as the computational backbone of this approach. C2S-scale is distinguished by its multimodal training paradigm^7^, in which transcriptomic profiles are encoded as “cell sentences” and jointly modeled with rich biological text and metadata, resulting in a corpus exceeding one billion tokens. With up to 27 billion parameters, the model demonstrates state-of-the-art performance across diverse tasks, including perturbation response prediction and multicellular reasoning. Importantly, prior work has shown that C2S-scale can generate experimentally validated, context-dependent predictions, including identifying conditional drug responses not explicitly represented in its training data^7^. Thus, this model provides empirical evidence for cross-context biological generalization, a critical requirement for virtual cell modeling.

Nevertheless, even broadly pretrained foundation models remain limited by the biological contexts represented in public datasets, particularly in clinically complex scenarios such as immune checkpoint inhibitor (ICI) therapy. This therapy involves dynamic, patient-specific processes, including heterogeneous CD8⁺ T cell states^8,9^, therapy-induced rewiring of effector programs^10^, and antigen-driven clonal expansion^11,12^, which are poorly captured in existing training corpora. To begin addressing this limitation, we incorporated paired single-cell RNA sequencing (scRNA-seq) and T-cell receptor sequencing (scTCR-seq) datasets from patients with advanced cancer^13,14^, sampled longitudinally at baseline and following ICI therapy^15^, and encompassing both responders and non-responders^13,14^.

Notably, specimen collection was dictated by predefined clinical monitoring schedules rather than experimental design; therefore, each sampling point likely represents a transient equilibrium state within the evolving temporal landscape of T-cell responses^16^ rather than a perfectly synchronized stage of immune dynamics. By integrating real-world clinical perturbation trajectories and clonotype-resolved dynamics, we captured key dimensions of T-cell response heterogeneity that are largely absent from existing training datasets, thereby fine-tuning the C2S-scale foundation model toward mechanistically grounded and clinically translatable predictions.

Building on this integrated framework, we further introduce a mathematically grounded representation of T cell functional dynamics that explicitly separates latent effector capacity from realized effector output^17,18^. This formulation allows the virtual cell model to move beyond static descriptions of cellular states and instead infer hidden functional identity and their transitions under therapeutic perturbation. Using this integrated AI-enabled virtual cell model, we identify a previously underappreciated latent effector state of CD8⁺ T cells in patients with advanced cancer, which is characterized by distinct signaling activity and constrained transcriptional output. We demonstrate that this state forms a functional reservoir that shapes the T cell response to ICI therapy, supporting a model in which ICIs primarily unmask pre-existing effector capacity rather than inducing de novo differentiation^17^. Together, this work establishes a generalizable strategy for integrating foundation models with mechanistic approach to construct interpretable virtual cell systems and provides new insight into the cellular and dynamical basis of ICI therapy for cancer.

## Results

### Analytical strategies for detection of latent effector state in virtual CD8⁺ T cells

To identify CD8⁺ T-cell functional states modulated by ICI therapy, we built upon our previously developed mathematical framework of PD-1 signaling^17^. In this framework, PD-1 (one of the key immune checkpoint molecules) functions as a reversible and graded masking mechanism that suppresses effector function without eliminating the underlying effector potential. The model provides a mechanistic explanation for several hallmark features of PD-1 blockade therapy, including preserved effector readiness in exhausted T cells, rapid functional restoration following checkpoint blockade, and heterogeneous clinical responses. Importantly, the theoretical model predicts that PD-1 blockade does not generate entirely new effector programs, but it reveals pre-existing and therapeutic reversible latent effector capacity ^17^.

Guided by this framework, we hypothesized that PD-1 blockade promotes the transition of CD8⁺ T cells from a latent effector state to an active effector state in patients with advanced cancer. To quantitatively represent this concept, we defined the latent effector state as a vector *E*(*t*) ∈ ℝ^*n*^, capturing a poised but submaximal effector program characterized by coordinated expression of canonical effector-associated genes, including *NKG7*, *CX3CR1*, *PRF1*, *KLRG1*, and *GZMB*. Unlike conventional transcriptomic measurements, *E*(*t*) represents latent functional capacity rather than observable transcriptional output. Thus, it encodes the potential of a T cell to rapidly acquire full effector function when inhibitory constraints are relieved during PD-1 blockade. To link latent effector capacity to observable gene expression, we modeled the dynamics of active effector-gene expression, *G*(*t*), using a first-order dynamical system described by ordinary differential equation:

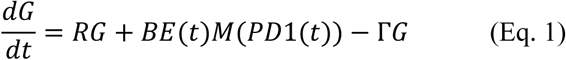

where *R* denotes the gene-regulatory matrix governing interactions among effector genes, *B* represents the conversion of latent effector capacity into active transcriptional output, and Γ describes gene-specific transcript turnover. The PD-1 masking function, *M*(*PD*1(*t*)), acts multiplicatively on the realization of latent effector capacity into measurable gene expression while leaving the latent state *E*(*t*) intact. Equation 1 provides a mechanistic representation of active effector-gene expression by integrating four key processes: gene-regulatory interactions, latent-state realization, PD-1-mediated masking, and transcript turnover. Within this framework, diminished effector-gene expression in exhausted T cells reflects transcriptional suppression rather than loss of effector identity. Effector potential remains intact but is constrained by checkpoint signaling^19,20^. Consequently, PD-1 blockade is predicted to restore effector-gene expression by releasing the masking constraint and enabling rapid realization of pre-existing latent capacity, without requiring extensive de novo cellular reprogramming. This formulation therefore establishes a quantitative bridge between latent cellular potential and measurable transcriptional output, providing a theoretical foundation for identifying hidden effector states in virtual CD8⁺ T cells and supporting a model in which ICI therapy primarily unmasks pre-existing effector capacity rather than inducing de novo differentiation^14,21–23^.

### C2S-derived axis modeling reveals a latent effector-like T cell state embedded within the CD8^+^ T cell continuum

To identify functional states within human peripheral blood CD8⁺ T cells, we used C2S-Scale as a pretrained representation model to convert ranked single-cell transcriptomes into a biologically informed latent space. Using anchor cells from Dataset 1 (DS1) and Dataset 2 (DS2), we then constructed a supervised effector-to-non-effector axis within this latent space. PHATE was used to visualize the resulting cellular manifold^24^, while PROGENy^25^ pathway analysis provided an independent pathway-level interpretation of C2S-defined states. Dataset 3 (DS3), which included paired T-cell receptor (TCR) clonotype information, was used as an independent validation cohort. The overall study design, C2S-Scale architecture, and analytical workflow are summarized in **Supplementary Fig. 1A–B**.

Projection of CD8⁺ T cells onto the C2S-derived effector-to-non-effector axis revealed a distinct transcriptional identity enriched for effector-associated genes but located outside the conventional terminal effector region of the continuum. We operationally designated cells within this region as Latent_effector cells, reflecting their predicted effector-like potential despite their noncanonical transcriptional position. To assess the robustness of this finding, we performed a series of sensitivity analyses using different datasets, response groups, axis intervals, marker-gene programs, and C2S-axis model configurations (**Supplementary Fig. 2**). Across all parameter settings, a Latent_effector population remained detectable, although its estimated frequency varied substantially. Broader latent-axis intervals and more inclusive exhaustion-oriented gene programs generally yielded higher estimated frequencies, whereas narrower intervals and more restrictive cytotoxic or terminal-effector gene programs produced lower estimates (**Supplementary Fig. 2; Supplementary Table 1**). Additional variation was observed across dimensionality settings, including PCA50-Ridge, PCA100-Ridge, and full-dimensional Ridge models. Collectively, these findings suggest that the abundance of Latent_effector cells is influenced by the functional composition of the underlying T-cell response rather than reflecting a fixed, discrete T-cell subset.

Importantly, despite variation in estimated frequency, the Latent_effector state was reproducibly detected across all analytical settings, supporting the existence of an underlying C2S-defined biological state. Therefore, we considered the continuous C2S effector-to-non-effector axis to be the primary quantitative output of the model. For downstream biological analyses, we used the less stringent PCA50-Ridge classification to assign cells as Effector, Latent_effector, NonEffector, or Other, while a more stringent PCA50-Ridge definition was retained as a sensitivity analysis to evaluate the robustness of conclusions.

To further evaluate the stability of the model, we performed leave-one-patient-out (LOO) analyses (**Supplementary Fig.3**). Across both DS1 and DS2, the continuous C2S effector-to-non-effector axis was highly preserved, with strong correlations between the full-model and LOO-derived axis scores. In contrast, LOO models consistently identified smaller numbers of Latent_effector cells and showed only modest overlap with full-model Latent_effector assignments at the individual-cell level. These results indicate that the continuous C2S-derived axis is highly stable across patients, whereas threshold-based classification of Latent_effector cells is more sensitive to model calibration and represents a conservative operational annotation. Taken together, these analyses support a model in which a latent effector-like state is embedded within the broader CD8⁺ T-cell functional continuum. Consequently, throughout the remainder of the study, we interpret the continuous C2S effector-to-non-effector axis as the principal quantitative measure of cellular state, while Latent_effector labels are used as operational annotations to facilitate biological interpretation.

### A latent effector-like state is consistently identified among peripheral blood CD8⁺ T cells in patients with advanced cancer

Despite the transformative clinical success of ICI therapy, most patients with advanced cancer do not achieve durable responses. A key challenge in cancer immunology is to identify the CD8⁺ T-cell states that drive effective antitumor immunity and therapeutic responses^10,11^. By analyzing peripheral blood CD8⁺ T cells from three independent cohorts of patients with advanced melanoma, we consistently identified a previously unrecognized Latent_effector cell (**Fig. 1**). This population occupied a distinct position along the C2S-derived effector-to-non-effector continuum and was clearly distinguishable from canonical Effector, Memory, and NonEffector states. The reproducible detection of this population across model-development and independent validation cohorts suggested the existence of a conserved effector-adjacent CD8⁺ T-cell state with potential relevance to antitumor activity and response to PD-1 blockade. In the C2S UMAP representation, Latent_effector cells did not form a discrete, isolated cluster. Instead, they were embedded within broader CD8⁺ T-cell transcriptional neighborhoods and were frequently located adjacent to or intermingled with Effector cells across all datasets. Because UMAP primarily preserves local relationships within the embedding space and is not designed to infer developmental trajectories, these observations were interpreted together with complementary analyses, including the continuous C2S axis, PHATE visualization, marker-gene expression, pathway activity profiling, and TCR clonotype analyses.

**Figure. 1.**
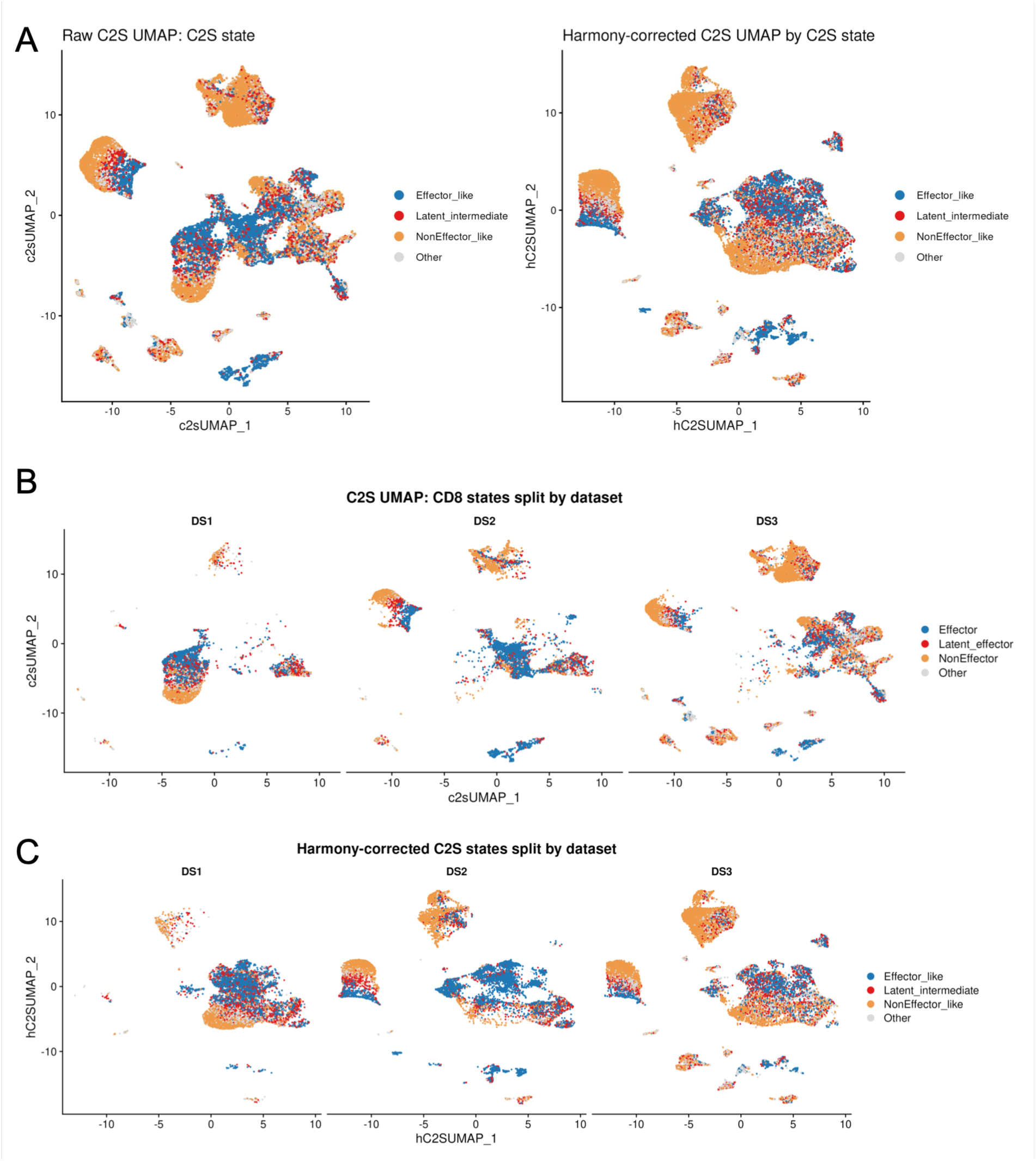
UMAP of C2S embeddings colored by relaxed state labels in CD8^+^ T cells among three datasets with or without Harmony correction. Three datasets (DS) were used in training (DS1 and DS2) and validation (DS3) of our integrative AI-enabled model based on C2S-scale. (A) Overall C2S UMAP used for visualizing a continuous semantic similarity in local C2S neighborhoods before and after Harmony correction. The clusters represent regions of similar transcriptional context. Latent effector cells were identified inside the largest clusters among a broader spectrum of CD8⁺ T cell state. (B-C) C2S UMAP split by dataset before (B) and after (C) Harmony correction.

To further define the relationship among Latent_effector, Effector, and NonEffector states, we visualized the C2S embedding using PHATE. PHATE was used to examine the overall organization of cells within the C2S-derived manifold, whereas the continuous C2S effector-to-non-effector axis was retained as the primary quantitative measure of CD8⁺ T-cell state. Within this framework, PHATE islands were interpreted as neighborhoods of cells sharing similar C2S embedding profiles and therefore similar ranked-gene-expression contexts captured by the C2S-Scale model, rather than as automatically defined cell types or developmental lineages. Visualization of the continuous C2S axis within PHATE space revealed a graded organization of CD8⁺ T-cell states across DS1, DS2, and DS3 (**Fig. 2A**). Cells occupying the same PHATE neighborhood often spanned a range of C2S axis values, particularly in DS3, indicating that the C2S manifold is not composed of strictly discrete cellular states. Rather, the C2S axis captured a continuous effector-to-non-effector spectrum across the peripheral CD8⁺ T-cell compartment. When relaxed C2S-derived state labels were overlaid onto the PHATE manifold, Latent_effector cells were readily observed in all three datasets but again did not form a fully isolated population (**Fig. 2B**). Instead, they occupied regions intermediate between canonical Effector and NonEffector states and were frequently embedded within larger CD8⁺ T-cell neighborhoods. This consistent spatial positioning supports the interpretation that the latent effector-like state represents a transitional or intermediate functional program situated along a continuum of CD8⁺ T-cell states rather than a separate cell type.

**Figure. 2.**
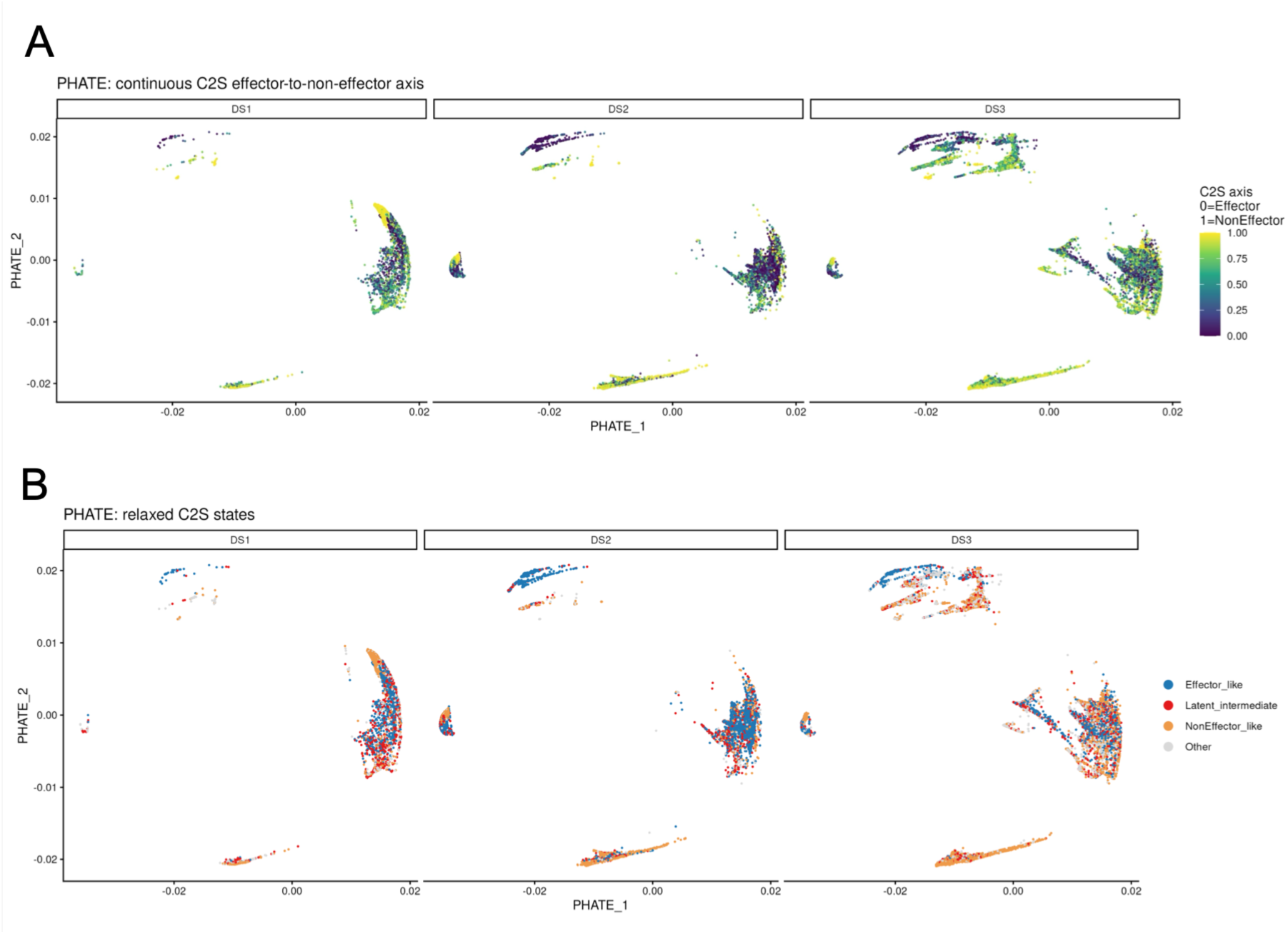
PHATE visualization split by dataset. (A) PHATE colored by the continuous C2S effector-to-non-effector axis score across datasets using the actual PHATE coordinate range and displays the C2S axis from 0 to 1, where 0 corresponds to Effector and 1 to NonEffector. (B) Thresholded categorical relaxed C2S states on the same PHATE manifold.

Marker-gene analysis further supported this interpretation (**Fig. 3**). Latent_effector cells retained expression of key cytotoxic effector-associated genes^14,26–28^, including CX3CR1, GZMB, NKG7, and PRF1, while simultaneously expressing genes commonly associated with progenitor/stem-like^29–31^ or exhaustion-related programs, including TCF7, IL7R, PDCD1, and TOX^32,33^. Thus, Latent_effector cells exhibit a hybrid transcriptional profile that combines core cytotoxic effector features with elements of progenitor and exhaustion-associated programs. Together, these findings suggest that Latent_effector cells represent an effector-like but partially restrained CD8⁺ T-cell state, positioned between canonical effector and non-effector states within the broader CD8⁺ T-cell functional continuum.

**Figure 3.**
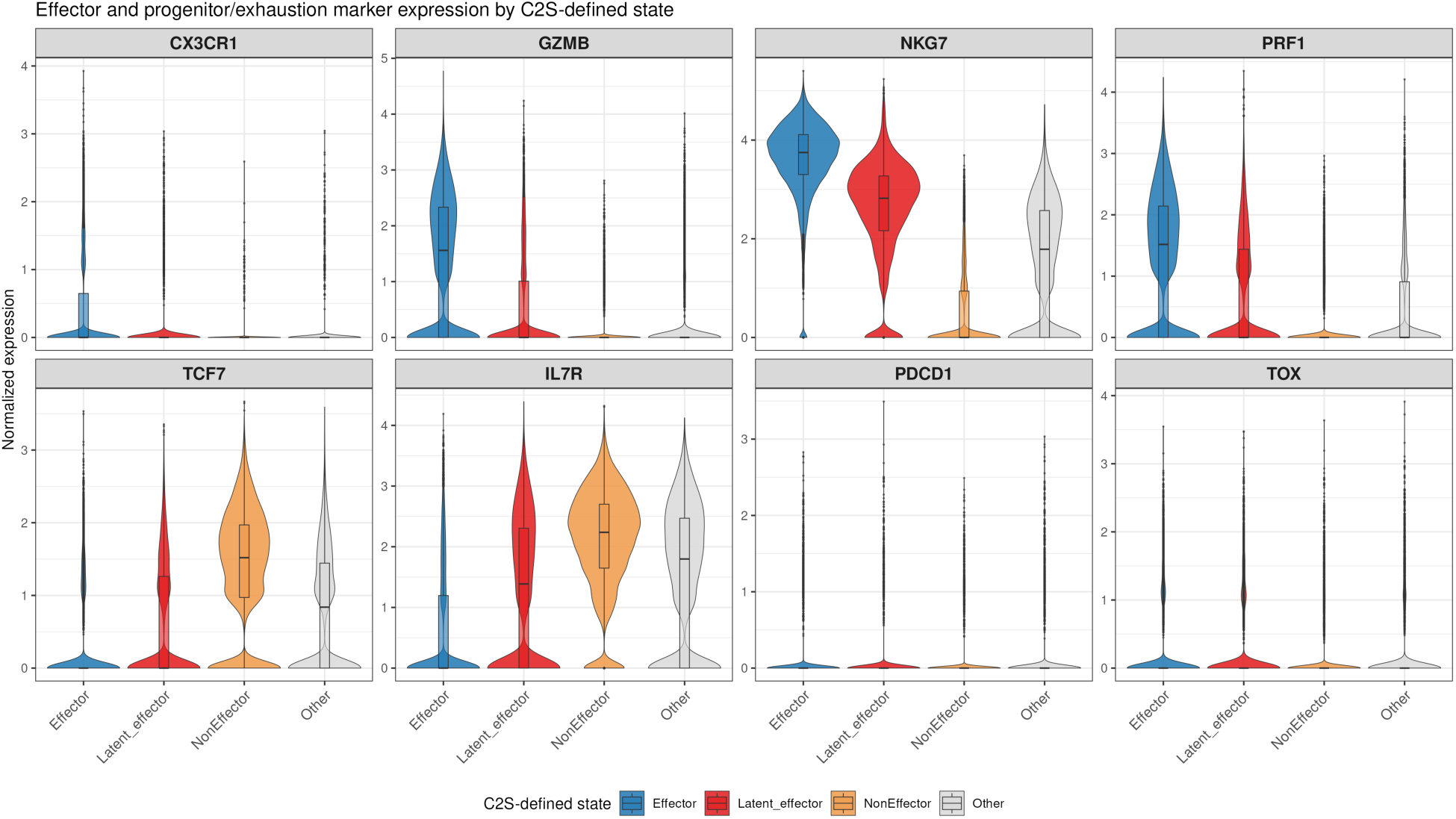
Gene expression of selected marker genes in C2S-defined T cell states. Effector genes: CX3CR1, GZMB, NKG7, PRF1. Progenitor/exhaustion-associated genes: TCF7, IL7R, PDCD1, TOX.

### Clonal enrichment and therapy-associated dynamics of the latent effector state during PD-1 blockade

Changes in T-cell receptor (TCR) clonotype size are a well-established feature of peripheral CD8⁺ T-cell responses to ICI therapy^23,34^, reflecting antigen-driven selection, expansion, and remodeling of the T-cell repertoire^21,30^. To determine how these antigen-driven responses are distributed across functional CD8⁺ T-cell states, we evaluated clonotype enrichment within the Effector, Latent_effector, NonEffector, and Other populations. This analysis allowed us to assess whether the latent effector state is associated with antigen-experienced T-cell clones and how this compartment changes during PD-1 blockade. Stratification of TCR clonotypes by clone size (large, medium, and small) demonstrated that both Effector and Latent_effector populations were significantly enriched for medium and large clones relative to other CD8⁺ T-cell states (**Fig. 4A–B**). Because clonally expanded effector T cells are typically generated through antigen-driven activation^12,20,35,36^, this enrichment suggests that Latent_effector cells, like canonical effector cells, arise from actively engaged and expanded T-cell clones^34^. Thus, antigen stimulation appears to preferentially populate both the Effector and Latent_effector compartments with cells possessing cytotoxic potential.

**Figure 4.**
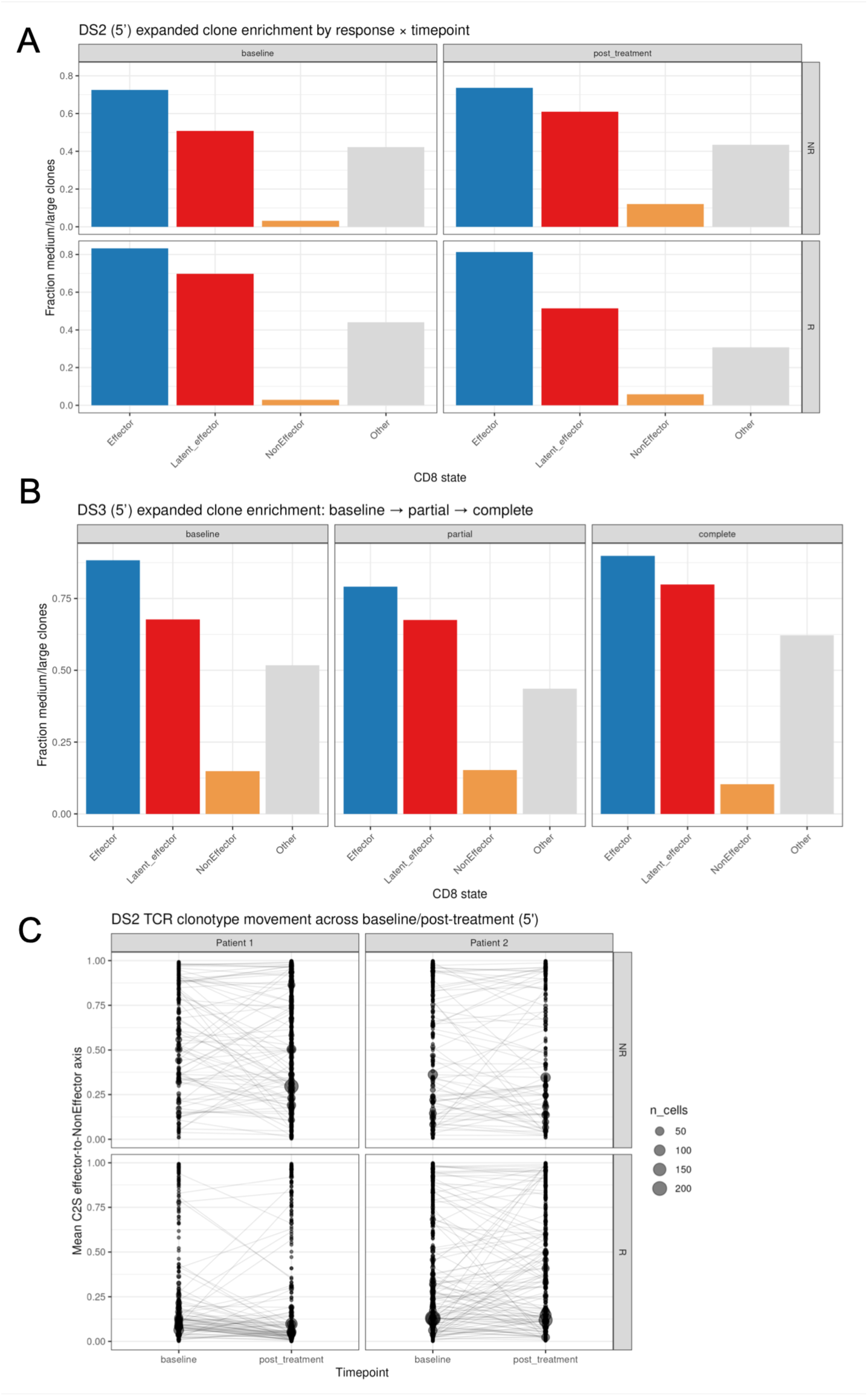
TCR clonality in latent state of CD8^+^ T cells. (A-B) Fraction of large and medium clones (clone size >= 3, treated as expanded) across C2S T cell states, in (A) DS2 by time points (baseline, post_treatment) and response (NR:Non-Responder, R:Responder), in (B) DS3 by time points (baseline, partial and complete response). (C) TCR clonotype movement across time points in DS2 by response. Lower mean C2S effector-to-NonEffector axis score is more effector-like.

To investigate the clinical relevance of this observation, we examined changes in the proportion of medium- and large-sized clonotypes within the Latent_effector compartment before and after PD-1 blockade in the DS2 cohort (**Fig. 4A**). In responders, the fraction of medium and large clones residing within the Latent_effector state decreased following therapy relative to baseline. In contrast, this metric remained largely unchanged in non-responders. One interpretation of this pattern is that, in responding patients, clonally expanded Latent_effector cells may undergo further differentiation or mobilization toward more overt effector states after PD-1 blockade^34^. Supporting the stability of the latent compartment itself, analysis of the independent validation cohort (DS3) showed persistent clonal enrichment of Latent_effector cells across baseline, partial-response, and complete-response samples (**Fig. 4B**), indicating that antigen-experienced clones can be maintained within this intermediate state over time.

We next asked whether clonal behavior could be detected along the continuous C2S effector-to-non-effector axis. Using paired baseline and post-treatment samples from DS2, we tracked individual TCR clonotypes and calculated the mean C2S axis score of each clone (**Fig. 4C**). In non-responders, clonotypes remained broadly distributed across the axis, with many clones occupying intermediate or non-effector regions following treatment. In contrast, clonotypes from responders were preferentially concentrated toward the effector end of the axis, consistent with increased occupancy of effector-like states after PD-1 blockade. These findings support the notion that successful therapy is associated with a shift of antigen-experienced T cell clones (not limited to tumor antigen-stimulated) toward a more effector-oriented functional state. As a complementary analysis, we quantified the proportion of cells from each clonotype that occupied the threshold-defined Latent_effector state (**Supplementary Fig. 4**). Individual clonotypes showed substantial variability in latent-state occupancy, indicating that clonally related cells can distribute across multiple functional states rather than remaining confined to a single phenotype. This observation further supports the view that the latent effector state represents one component of a broader continuum of CD8⁺ T-cell differentiation.

Collectively, these findings demonstrate that Latent_effector cells are enriched for clonally expanded, antigen-experienced T cells and undergo therapy-associated changes during PD-1 blockade. The concordance between TCR clonotype dynamics and the C2S-derived functional landscape provides independent evidence that the Latent_effector state represents a biologically meaningful effector-adjacent program rather than a transcriptional artifact. Moreover, the preferential movement of responder clonotypes toward the effector end of the continuous C2S axis suggests that the latent effector compartment may serve as a reservoir of antigen-experienced cells with the capacity to contribute to productive antitumor immune responses.

### Tumor-specific CD8⁺ T cells are preserved within the latent effector state in patients responding to PD-1 blockade

To determine whether the latent effector compartment contains tumor-reactive CD8⁺ T cells that can contribute to successful immunotherapy responses, we longitudinally tracked tumor antigen-specific T-cell clonotypes in peripheral blood from a patient in the DS3 cohort who achieved a clinical response to PD-1 blockade therapy. Samples were collected at baseline before treatment, during therapy, and at the time of complete clinical response. Using 10x Genomics BEAM-T technology, we identified a TCR clonotype specific for the cancer-testis antigen NY-ESO-1 (Ag-12-Com) and followed its distribution across C2S-defined functional states over time (**Fig. 5**). At baseline, the NY-ESO-1-specific clonotype was predominantly detected within the Latent_effector compartment rather than within the canonical Effector population, indicating that tumor-reactive cells can exist in an effector-adjacent but functionally restrained state before therapy. During treatment, this clone exhibited dynamic changes, including a transient decline during the partial-response phase, followed by marked expansion coincident with complete clinical response. The emergence of a large population of tumor-specific cells after PD-1 blockade suggests that pre-existing antigen-experienced clones retained the capacity for expansion and functional engagement once inhibitory constraints were relieved. Although derived from a single patient, this longitudinal observation provides proof-of-concept evidence that tumor-specific CD8⁺ T-cell clones can be maintained within the latent effector state prior to treatment and subsequently expand during successful PD-1 blockade. These findings support a model in which the latent effector compartment serves as a reservoir of clonally competent, tumor-reactive T cells that can contribute to effective antitumor immunity following immune checkpoint inhibition.

**Figure 5.**
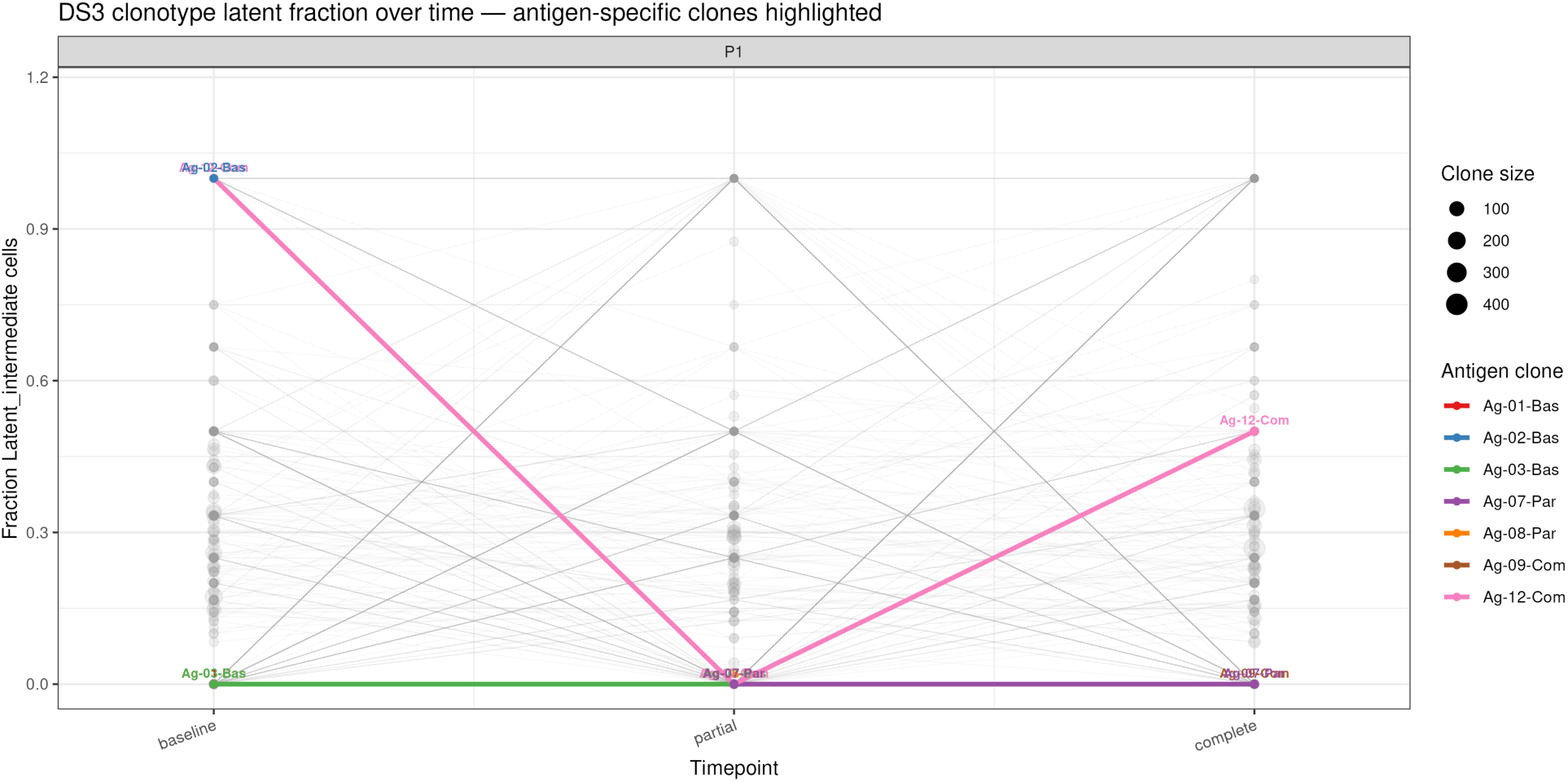
Tracking tumor antigen-specific TCR clones over time of clinical responses to anti-PD-1 therapy. Tumor antigen (NY-ESO-1) specific TCR clones were identified using BEAM-T method for scTCR/scRNA-seq in DS3 overtime from baseline through partial and complete clinical response to anti-PD-1 therapy. Each line represents each TCR clones and changes overtime in fraction of latent effector state.

Notably, these biological observations are consistent with the predictions of our mathematical framework^17^. While Eq. 1 describes the dynamics of the active effector program, *G*(*t*), we introduced a second variable, *Q*(*t*), to represent the cumulative functional output generated by CD8⁺ T cells over time. Unlike *G*(*t*), which reflects the instantaneous activity of an effector-associated gene-expression program, *Q*(*t*) represents the accumulated biological consequences of that activity, such as sustained cytotoxic function and antitumor activity. We therefore define cumulative functional output as:

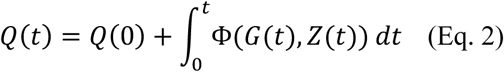

where *Q*(0) denotes the baseline functional output before therapy. The variable *Z*(*t*) represents context-dependent factors that influence how molecular activity is translated into measurable T cell functionality, including antigen availability, immune-synapse formation, target-cell accessibility, and other extrinsic regulatory constraints. The function Φ therefore serves as a phenomenological link between intracellular molecular states and observable immune function. Of note, Eq. 2 does not represent a simple integration of Eq. 1, but defines a downstream functional output generated by the effector program described by *G*(*t*). Thus, two cells with similar levels of effector-associated gene expression may differ substantially in realized functional output depending on the biological context captured by *Z*(*t*). Within this framework, the rapid clinical and immunological responses observed after PD-1 blockade can be explained by the release of pre-existing effector potential rather than by the generation of entirely new effector populations de novo^13,23,37^.

The longitudinal behavior of the NY-ESO-1-specific clonotype is particularly consistent with this model. Before therapy, this tumor-reactive clonotype resided predominantly within the Latent_effector compartment, indicating that it had already acquired features of an effector program but remained incompletely expressed at the functional level. Following PD-1 blockade, the same clonotype underwent marked expansion coincident with clinical response, consistent with the mobilization of pre-existing effector potential and increased contribution to cumulative functional output. These observations support the existence of a reservoir-like state in which tumor-reactive cells retain effector capacity while remaining subject to inhibitory regulation.

Analysis of the broader peripheral CD8⁺ T-cell repertoire further reinforced this interpretation. Although many clonotypes likely recognize non-tumor antigens, including pathogen-derived antigens, medium- and large-sized clones were preferentially enriched within both the Latent_effector and Effector compartments (**Fig. 5**, gray lines and dots). This finding indicates that antigen-experienced T cells are not confined to terminal effector states. The substantial populations of expanded clonotypes can persist within the Latent_effector compartment while retaining the capacity for subsequent activation, differentiation, or functional output. Thus, TCR clonal architecture provides an independent line of evidence supporting the biological relevance of the C2S-defined latent effector state. Taken together, these findings connect tumor specificity, clonal expansion, and therapeutic response to the latent effector compartment and provide experimental support for the central predictions of Eq. 1 and Eq. 2. The identification of tumor-reactive clonotypes within the Latent_effector state therefore provides mechanistic support for the concept that this compartment serves as a clinically relevant reservoir of antitumor immunity whose functional contribution accumulates over time as reflected by *Q*(*t*).

### Signaling pathway activity reveals that the latent effector state is signaling-active but functionally restrained

The Latent_effector state was identified primarily from its position along the C2S-derived effector-to- non-effector continuum rather than from a conventional transcriptional signature. Because this state is hypothesized to represent latent effector capacity that is not fully translated into overt effector function, standard gene-expression–based pathway analyses may underestimate its biological activity^25^. Conventional pathway enrichment methods typically depend on the expression levels of pathway member genes and therefore may fail to capture pathway engagement when downstream transcriptional outputs remain partially suppressed, as predicted by our PD-1 masking model and mathematical framework^17^. To address this limitation, we applied PROGENy^25^, a pathway-inference approach that estimates signaling activity from downstream perturbation-responsive transcriptional footprints rather than from the expression of pathway genes themselves. This strategy provides an orthogonal view of cellular signaling and is particularly useful for identifying pathways that may be functionally active even when classical transcriptional signatures are incomplete. PROGENy scores were calculated from the normalized single-cell gene-expression matrix and used to characterize signaling programs associated with the C2S-defined Effector, Latent_effector, and NonEffector states.

We first evaluated pathway activity at the single-cell level and subsequently aggregated pathway scores in a pseudobulk-like manner by dataset, treatment timepoint, response status, and C2S-defined state. This dual approach allowed us to determine whether signaling activity could further distinguish the latent effector state from canonical Effector and NonEffector populations and provide additional biological insight into its functional properties. Remarkably, PROGENy analysis revealed well-defined and reproducible signaling profiles across the three major CD8⁺ T-cell states, with relatively little overlap between T cell states and groups (**Fig. 6**). In several datasets, pathway-level separation was even more pronounced than separation based on transcriptional features alone, suggesting that signaling activity provides complementary information that is not fully captured by gene-expression patterns.

**Figure 6.**
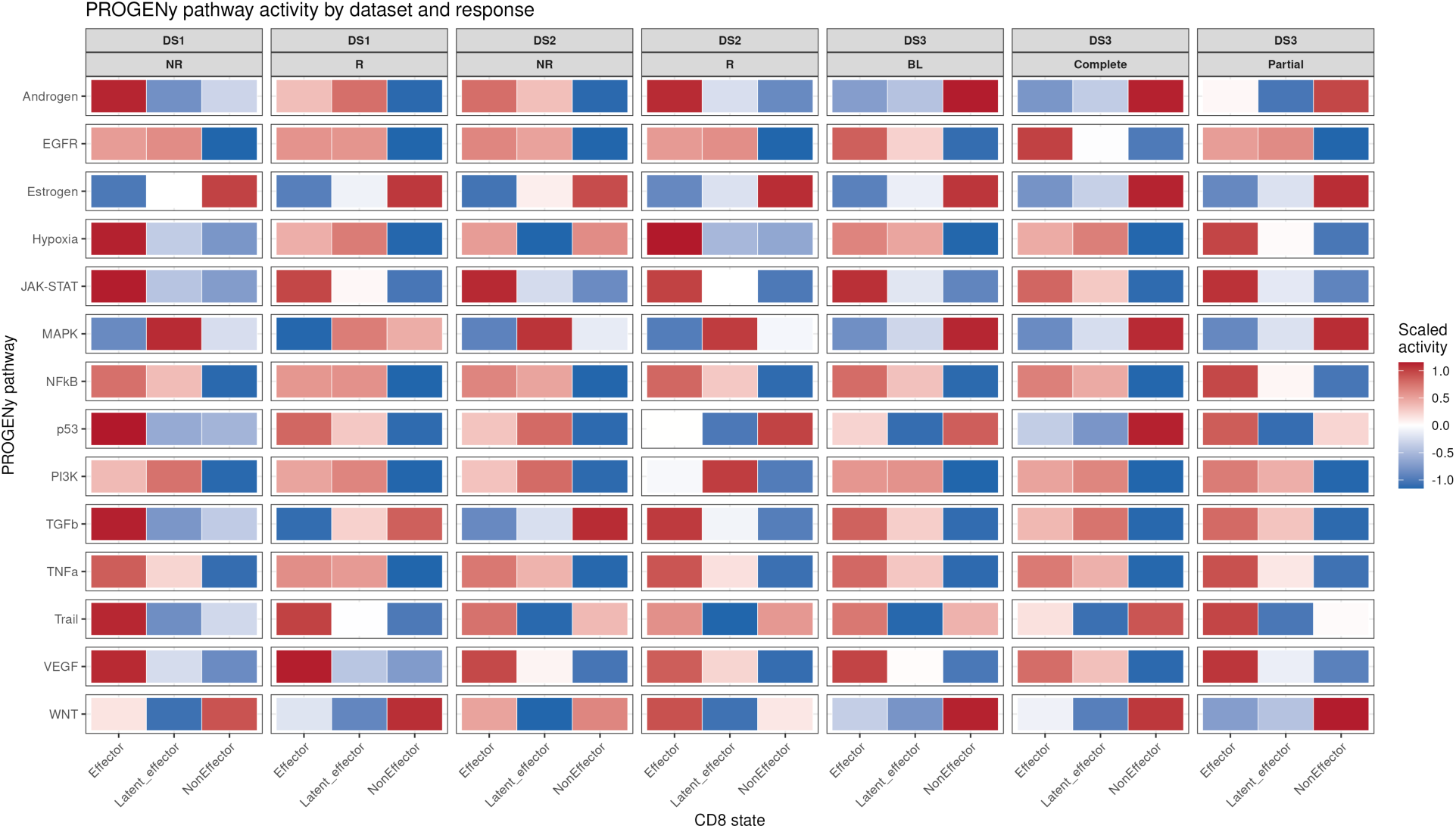
PROGENy pathway activity across C2S cell states split by dataset. Colors indicate scaled PROGENy pathway activity from high to low (1.0 to −1.0), showing relative differences within pathway/dataset/response comparisons among clinical responses to anti-PD-1 therapy I DS1 and DS2, and DS3 with timeline of responses. NR: Non-responders, R: responders; BL: baseline; Complete: complete responsive stage; Partial: partial responsive stage.

Across datasets, Latent_effector cells consistently occupied an effector-adjacent signaling state. Compared with NonEffector cells, Latent_effector cells displayed increased activity in inflammatory and cytokine-responsive pathways, including TNFα/NF-κB and JAK-STAT signaling. However, pathway activity generally remained lower than that observed in canonical Effector cells. This intermediate signaling profile is consistent with a state that retains substantial effector-associated signaling potential while not fully achieving the signaling output characteristic of terminal effector function. Additional pathways supported a similar interpretation. MAPK signaling showed evidence of increased activity in Latent_effector cells in DS1 and DS2, although the magnitude of this effect varied across datasets and treatment timepoints. In contrast, PI3K pathway activity^38–40^ was relatively preserved across multiple datasets and response groups, suggesting that Latent_effector cells maintain signaling competence and retain the capacity to respond to activating stimuli. By comparison, we found the pathways including WNT, p53, TGFβ, hypoxia, androgen, and estrogen signaling demonstrated substantial context-dependent variability and did not consistently distinguish the Latent_effector state from other CD8⁺ T-cell populations. Consequently, these pathways were not interpreted as defining features of the latent effector program. Taken together, these findings indicate that Latent_effector cells are not biologically quiescent but instead preserve signaling programs typically associated with activated and responsive T cells.

Overall, the PROGENy results provide an important functional dimension to the interpretation of the C2S-defined latent effector state. The PROGENy findings suggest that many Latent_effector cells have already activated key inflammatory and cytokine-response signaling pathways but have not yet fully translated this signaling activity into the complete effector transcriptional and functional program. Thus, the latent effector state is best viewed as signaling-active and effector-adjacent, yet functionally restrained, providing a plausible cellular substrate for the rapid reinvigoration of antitumor immunity following PD-1 blockade.

### Latent Effector Cells as a Probabilistic Biomarker of Response to Immune Checkpoint Inhibitor Therapy

To evaluate the clinical relevance of the C2S-defined Latent_effector state, we analyzed an independent clinical cohort of patients with advanced melanoma treated with immune checkpoint inhibitors (ICIs), either alone or in combination with chemotherapy14 (**Table 1**). In our previous study, we and others have identified an expansion of peripheral blood CX3CR1⁺GZMB⁺ CD8⁺ T cells in patients who responded to PD-1 blockade therapy for advanced melanoma or lung cancer^14,22,41,42^. These cells exhibited an effector-memory (Tem)–like phenotype and were strongly associated with clinical benefit. Notably, CX3CR1 has been widely recognized as a marker of antigen-experienced CD8⁺ T cells undergoing differentiation from stem-like or progenitor-like states toward terminal effector or exhausted states in both preclinical and clinical settings^43,44^. Therefore, based on their phenotype and functional characteristics, we considered circulating CX3CR1⁺GZMB⁺ CD8⁺ T cells as a clinically measurable surrogate for the C2S-defined Latent_effector population predicted by our mathematical framework and single-cell analyses.

**Table 1.**
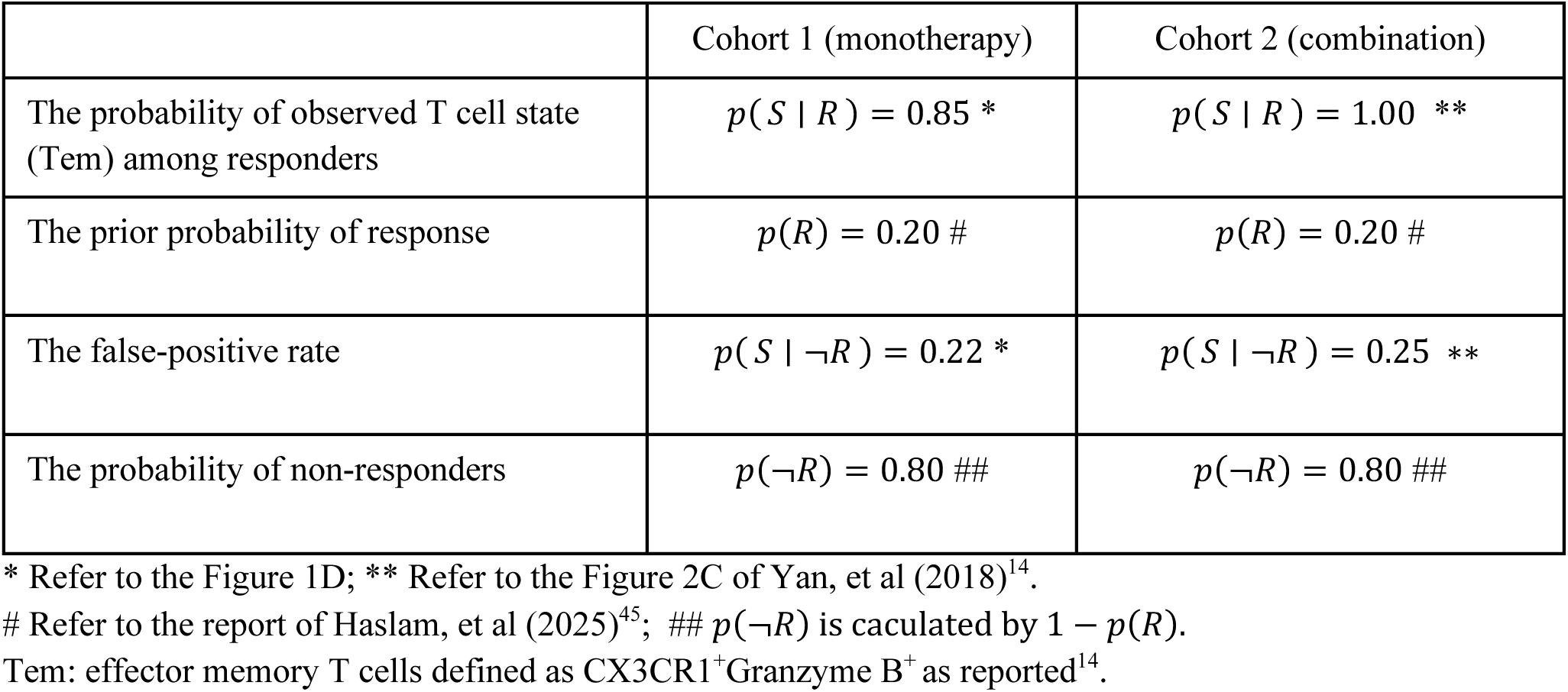
Estimation of the probability of clinical responses to ICI monotherapy or combination therapy.

To quantitatively assess the relationship between this T-cell state and therapeutic response, we applied a Bayesian framework to estimate the probability that a patient is in a responder state (*R*) given the observation of a predefined T-cell state (*S*). Here, *S* was defined as the presence of CX3CR1⁺GZMB⁺ CD8⁺ T cells above a threshold frequency previously identified as being associated with anti-PD-1 responsiveness^14^. In the first clinical cohort (Cohort 1), which consisted of patients treated with ICI monotherapy (**Table 1**), the probability of observing this T-cell state among responders, *P*(*S* ∣ *R*), was 0.85. The prior probability of response, *P*(*R*), was set at 0.20, consistent with the approximate overall response rate reported for advanced melanoma patients receiving ICI therapy^45^. Among non-responders, the probability of observing the same T-cell state, *P*(*S* ∣ ¬*R*), was 0.22, yielding a non-responder prior probability *P*(¬*R*) = 0.80.

Using the law of total probability,

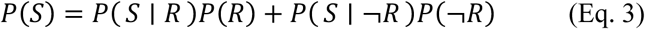

the overall probability of observing the signature was calculated as approximately 0.35. Application of Bayes’ theorem then yielded:

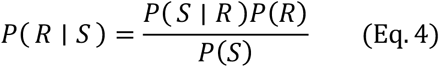

resulting in a posterior probability of response of approximately 48.6%. Thus, the presence of the CX3CR1⁺GZMB⁺ CD8⁺ T-cell signature increased the predicted likelihood of response from the baseline population rate of 20% to nearly 50%, more than doubling the probability of clinical benefit. We next evaluated an independent cohort of patients treated with combination ICI plus chemotherapy (**Table 1**). In this cohort, the likelihood of observing the signature among responders, *P*(*S* ∣ *R*), approached 1.0, whereas the false-positive rate among non-responders, *P*(*S* ∣ ¬*R*), was 0.25. Using the same prior response probability (*P*(*R*) = 0.20), Bayesian analysis yielded a posterior response probability of approximately 50%, remarkably similar to that (48.6%) observed in the monotherapy cohort as above. The reproducibility of these estimates across distinct therapeutic settings suggests that this effector-memory–like CD8⁺ T-cell state represents a robust, treatment-independent indicator of responsiveness to immune checkpoint blockade. Similar associations between circulating effector-like CD8⁺ T-cell populations and favorable clinical outcomes have been reported in multiple independent studies^22,41–44,46,47^.

These real-world clinical observations provide another independent validation of the biological relevance of the C2S-defined Latent_effector state. Phenotypically, these cells express cytotoxic effector markers, including CX3CR1 and GZMB, yet remain distinct from fully differentiated terminal effector cells. Instead, the combined clinical, transcriptional, and TCR clonotype data support a model in which Latent_effector cells occupy a transitional state along the differentiation continuum between stem-like/progenitor and terminal effector CD8⁺ T cells. Within this framework, PD-1 blockade does not primarily generate entirely new antitumor responses. Rather, it promotes the activation, expansion, and functional deployment of a pre-existing reservoir of antigen-experienced CD8⁺ T cells with latent effector potential. Consistent with this interpretation, enrichment of CX3CR1⁺GZMB⁺ CD8⁺ T cells above a critical threshold was associated with improved clinical outcomes, suggesting that the abundance of Latent_effector cells may serve as a clinically actionable biomarker for predicting responsiveness to PD-1 blockade therapy.

Taken together, the convergence of mathematical modeling, single-cell transcriptomics, TCR clonotype analyses, and independent clinical validation supports the existence of a biologically and clinically meaningful Latent_effector compartment composed of antigen-experienced CX3CR1⁺GZMB⁺ CD8⁺ T cells that retain substantial effector potential despite incomplete functional realization. Expansion or mobilization of this compartment following PD-1 blockade appears to be associated with therapeutic benefit, highlighting Latent_effector cells as both a mechanistically informative CD8⁺ T-cell state and a promising biomarker for patient stratification during ICI therapy.

## Discussion

In this study, we combined mathematical modeling, single-cell transcriptomics, TCR clonotype analysis, and a pretrained C2S representation model to identify a previously unrecognized latent effector-like state within the peripheral CD8⁺ T-cell compartment of patients with advanced melanoma. Across independent cohorts, this state was consistently positioned between canonical Effector and NonEffector populations along a continuous C2S-derived effector-to-non-effector axis. Multiple independent lines of evidence, including transcriptional profiling, pathway activity, clonal architecture, longitudinal treatment dynamics, and tumor antigen-specific T-cell tracking, support the existence of a biologically meaningful and functionally poised CD8⁺ T-cell state that is distinct from classical effector, memory, and exhausted populations.

A central finding of this work is that the latent effector state appears to represent a reservoir of antigen-experienced CD8⁺ T cells that retain substantial effector potential despite incomplete realization of effector function. This conclusion is supported by several observations. First, Latent_effector cells consistently expressed cytotoxic effector-associated genes, including CX3CR1, GZMB, NKG7, and PRF1, while also retaining features associated with progenitor-like^29,30,48^ or exhaustion-related programs^49–52^. Second, Latent_effector cells were enriched for medium and large TCR clonotypes, indicating prior antigen-driven expansion and active participation in adaptive immune responses^21,34^. Third, tumor antigen-specific CD8⁺ T-cell clones were detected within the latent effector compartment before treatment and subsequently expanded during successful PD-1 blockade. Collectively, these findings suggest that the latent effector state is not a transcriptional artifact but rather represents a biologically relevant intermediate state within the broader CD8⁺ T-cell continuum.

Our findings further support a model in which PD-1 blockade predominantly acts by releasing pre-existing effector potential rather than generating entirely new antitumor responses de novo^20,30,43^. This interpretation is consistent with our mathematical framework^17^, which distinguishes latent effector capacity from realized effector output. Within this framework, checkpoint signaling constrains the conversion of latent effector potential into active effector function, whereas PD-1 blockade alleviates this constraint and permits rapid functional deployment of pre-existing antigen-experienced T cells. The rapid expansion of tumor-reactive clonotypes observed following therapy, together with the persistence of clonally expanded cells within the latent compartment, provides biological support for this model and offers a potential explanation for the rapid response kinetics often observed clinically after PD-1 inhibition.

An additional insight from this study is that the latent effector state is characterized not only by its transcriptional profile but also by a distinct signaling landscape. Using PROGENy pathway inference, we found that Latent_effector cells retained activation of inflammatory and cytokine-responsive pathways, including TNFα/NF-κB and JAK-STAT signaling, despite failing to fully acquire the pathway activity profile of canonical effector cells. These findings indicate that Latent_effector cells are not quiescent or inactive. Rather, they represent a signaling-competent population that remains partially restrained from achieving full effector output. This distinction may explain why conventional transcriptome-based analyses alone have not consistently identified such cells and highlights the value of integrating pathway activity with transcriptional and clonal information.

Several conceptual advances emerge from this work. First, our framework emphasizes that CD8⁺ T-cell states are best viewed as positions along a continuous functional landscape rather than as strictly discrete cellular categories. Second, by separating latent capacity from realized effector function, our model provides a quantitative framework for interpreting checkpoint regulation as a reversible constraint on effector activity. Third, the integration of transcriptional, signaling, and clonotype information enables identification of biologically meaningful T-cell states that may not be apparent from conventional clustering approaches alone.

These findings also have important clinical implications. The presence of a latent effector reservoir may help explain why some patients achieve durable responses to PD-1 blockade whereas others do not. Patients harboring larger populations of clonally expanded Latent_effector cells may possess a greater pool of recruitable antitumor T cells capable of responding to checkpoint inhibition. Consistent with this possibility, a clinically measurable surrogate population of CX3CR1⁺GZMB⁺ CD8⁺ T cells was associated with therapeutic responsiveness across independent clinical cohorts^14,22,41^. Bayesian analysis demonstrated that the presence of this signature increased the likelihood of response from approximately 20% at baseline to nearly 50%, supporting its potential utility as a predictive biomarker. More broadly, these observations suggest that effective immunotherapy may depend not only on the abundance of terminal effector cells but also on the availability and mobilization of latent effector capacity from a transition phenotype (such as CX3CR1) of CD8^+^ T cells^28,43,44,53^.

Several limitations should be acknowledged. First, our conclusions are derived primarily from observational single-cell datasets and therefore do not provide direct experimental proof of state transitions between latent effector and effector populations. Definitive validation will require lineage-tracing approaches and prospective longitudinal studies with higher temporal resolution. Second, the latent effector state is defined computationally using the C2S-derived axis and associated analytical frameworks. Although supported by multiple independent lines of evidence, the precise molecular boundaries of this state remain to be established through functional experimentation. Third, our analyses were performed predominantly in peripheral blood samples from patients with melanoma, and it remains unclear whether analogous latent effector populations exist within tumor-infiltrating lymphocytes or across other cancer types. Finally, the mathematical model intentionally simplifies the complexity of T-cell regulation and does not explicitly incorporate additional factors such as metabolic programming, epigenetic regulation, or interactions with other immune cell populations. Importantly, this model does not incorporate the temporal oscillations of many regulatory pathways underlying T-cell function^16^.

Therefore, peripheral blood T-cell states may represent dynamic equilibria that transition between distinct regulated steady states, rather than stable, discrete phenotypes, and thus may not be accurately characterized by single or limited sampling at empirical time points.

Future studies should focus on experimentally validating the functional properties of Latent_effector cells and defining the molecular mechanisms that maintain this state. Integration of longitudinal single-cell multi-omics approaches, including scRNA-seq, scATAC-seq, and TCR lineage tracking, will be particularly valuable for determining whether Latent_effector cells possess epigenetic features consistent with rapid effector reactivation. Extending these analyses to tumor-resident T cells and to additional immunotherapy settings will help establish the generalizability of the latent effector concept. Furthermore, therapeutic strategies aimed at preserving, expanding, or mobilizing latent effector capacity may represent a promising avenue for enhancing the efficacy of checkpoint blockade.

In conclusion, our study identifies a latent effector-like CD8⁺ T-cell state that is antigen-experienced, clonally expanded, signaling-active, and poised for functional reactivation. This state bridges classical effector and non-effector populations within a continuous CD8⁺ T-cell landscape and provides a mechanistic framework linking T-cell state, checkpoint regulation, and clinical response. By integrating mathematical modeling with single-cell and clinical data, these findings support the concept that successful PD-1 blockade acts primarily by mobilizing pre-existing effector potential. Quantification of the latent effector compartment may therefore provide both mechanistic insight into antitumor immunity and a foundation for future biomarker development and patient stratification strategies in cancer immunotherapy.

## Methods

### Patient information and processing of peripheral blood samples

Peripheral blood samples for this study were collected after written consent was obtained from each participant. Peripheral blood leukocytes (PBMC’s) were isolated from patient blood samples via centrifugation with lymphoprep (07851, StemCell Technologies) and SepMate conical tubes (85450, StemCell Technologies). Samples were then stored in liquid nitrogen for later use and thawed on the day of experiments. Human CD8^+^ T Cell Enrichment Kit (StemCell Technologies) according to the manufacturer’s protocol. CD8^+^ T cells were then immediately stained with BEAM-peptide-MHC assemblies. Clinical course, treatment information, and outcomes in patients treated with anti-PD-1/L1 therapy at Mayo Clinic were retrospectively collected. Response to treatment was evaluated according to standard clinical practice guidelines using RECIST, where R=responders; NR=non-responders are noted using RECIST at the 12-week post-initiation of therapy time-point for all patients. The study was approved by the Mayo Clinic Rochester IRB and was conducted according to Declaration of Helsinki principles.

### Preparation and staining with BEAM-peptide-MHC assemblies

A tumor antigen peptide (NY-ESO-1) and a negative control peptide (10X Genomics) were mixed with 2.4 uL of BEAM conjugate (Chromium, Single Cell 5’ BEAM Core Kit PE, PN-1000539, 10X Genomics) and 3 uL of human MHC monomer (Chromium Human MHC Class I A0201 Monomer Kit, PN-1000542, 10X Genomics; Chromium Human MHC Class I A1101 Monomer Kit, PN-1000543, 10X Genomics) to prepare BEAM-conjugated, target peptide-versus negative control peptide-bound MHC assemblies, respectively. They were then incubated in dark for 30 minutes, quenched with 15 uL of quenching agent (10X Genomics), and pooled together for CD8^+^ T cell staining, during which purified CD8^+^ T cells (1 x 10^6^) were suspended in 90 uL of chilled PBS with 2% FBS and mixed with BEAM-peptide-MHC assemblies in dark for 15 minutes on ice. Samples were washed with 3.5 mL of chilled PBS with 2% FBS and centrifuged at 300 RCF for 5 minutes at 4 C° to remove free BEAM-peptide-MHC assemblies.

### Single-cell RNA and TCR sequencing

Single-cell RNA and TCR sequencing was performed by the Genome Analysis Core at Mayo Clinic. Cells were counted and measured for viability using a Vi-Cell BLU Cell Viability Analyzer (Beckman-Coulter). Barcoded Gel Beads were thawed from −80 C° and a cDNA master mix was prepared according to the manufacture’s instruction for Chromium Next GEM Single Cell 5’ Dual Index Kit v2 (10X Genomics). Based on the desired number of cells to be captured for each sample, a volume of live cells was mixed with the cDNA master mix. Of note, a per sample concentration of 500,000 to 700,000 cells per milliliter or better is required for the standard targeted cell recovery of 10,000 cells. Cell suspension and master mix, thawed Gel Beads, and partitioning oil were added to a Chromium Single Cell G chip. The filled chip was loaded into a Chromium X instrument, where each sample was processed and the individual cells within the sample were partitioned into uniquely labeled Gel Beads-In-Emulsion (GEMs). The GEMs were collected from the chip and taken to the bench for reverse transcription, GEM dissolution, and cDNA clean-up. The cleaned cDNA was amplified and size selected. The resulting cDNA, a pool of uniquely barcoded molecules, was used to generate 5’ gene expression (GEX) libraries and enriched TCR libraries. In addition, the supernatant from the cDNA clean-up step contained amplified BEAM DNA. During all library construction, standard Illumina sequencing primers and a unique i7 Sample index were added to each cDNA and DNA pool. The Dual Index TT Set A plate was used for GEX and TCR libraries; and the Dual Index TS Set A plate was used for BEAM libraries (10x Genomics). All cDNA pools and resulting libraries were measured using Qubit High Sensitivity assays (Thermo Fisher Scientific) and Agilent Bioanalyzer High Sensitivity chips (Agilent). The minimum acceptable amount of cleaned cDNA to use in library construction is 1 ng. The minimum amount of finished library that can be sequenced is 3nM. Gene expression libraries were sequenced at 50,000 read pairs per cell; TCR libraries were sequenced at 10,000 read pairs per cell; and BEAM libraries were sequenced at 10,000 read pairs per cell. Sequencing steps followed the standard Illumina protocol for the Illumina NovaSeq X Plus™. For all libraries, the flow cell was sequenced as paired-end 100 (x2) reads, using NovaSeq X Plus sequencing reagents and software.

### Data Preprocessing and Quality Control

The Dataset 1 and Dataset 2 have been published^13^ and were used as training dataset for this study, and the Dataset 3 (generated from BEAM-T approach) was used for independent validation. Raw 10x Genomics gene-expression and 5′ TCR sequencing data were processed with Cell Ranger v7.1.0 using the GRCh38-2020-A reference. Filtered feature-barcode matrices and, when available, filtered V(D)J contig-annotation files were used for downstream analysis. Filtered Cell Ranger matrices were imported into Seurat. Sample-level metadata, including dataset, patient identifier, time point, treatment response, chemistry, and TCR availability, were assigned from manually curated sample manifests. Cells were filtered using standard quality-control metrics, including detected genes, total UMI counts, and mitochondrial transcript fraction. In the current pipeline, cells were retained using thresholds of nFeature_RNA > 300, nFeature_RNA < 6000, nCount_RNA < 30000, and percent.mt < 20. Potential non-CD8 or contaminating populations were evaluated using marker-based UMAP, violin, and dot-plot inspection. Conventional Seurat clustering was used for QC and reference annotation, but not as the primary basis for defining the latent effector-like state. After QC, gene-expression data were log-normalized, variable genes were identified, and scaled data were generated with regression of mitochondrial percentage and total UMI count. PCA, UMAP, neighborhood graph construction, and Seurat clustering were performed to inspect sample structure and cell-state composition. These conventional analyses served as baseline reference analyses and quality checks for the C2S-based model. For samples with TCR sequencing, filtered contig-annotation files were processed with scRepertoire and linked to the corresponding Seurat barcodes. Productive TRA and TRB chains were used to assign paired clonotypes, including amino-acid clonotype identifiers (CTaa), TRA/TRB CDR3 sequences, clone-size categories, and clone-frequency summaries. TCR information was retained as an orthogonal validation layer and was not used in the primary C2S embedding step. This design allowed paired clonotype tracking, clone expansion and BEAM-T antigen-associated clone assessment to be evaluated independently of transcriptomic state assignment.

### Model Architecture Development

For C2S modeling, each cell was converted into a gene-only ranked sentence. Genes with nonzero normalized RNA expression were ranked in descending order within each cell, and the top-ranked (up to 1000) genes were concatenated into a space-delimited sentence. These gene-only sentences were passed to the pretrained C2S-Scale model (C2S-Scale-Pythia-1B) to generate one fixed-length embedding vector per cell. TCR-enhanced sentences were exported for secondary analyses, but the primary C2S embeddings were generated from gene-expression sentences only. The downstream C2S model was designed to convert gene-only C2S embeddings into an interpretable continuous CD8⁺ T cell effector-to-non-effector axis. High-confidence effector and non-effector/progenitor-exhaustion anchor cells were selected from the DS1 and DS2 discovery cohorts using the primary current_default marker program. Effector anchors were defined using cytotoxic and effector-associated genes, including GZMB, PRF1, GNLY, NKG7, IFNG, CX3CR1, FGFBP2, KLRG1, CCL5, and GZMA. Non-effector/exhaustion anchors were defined using checkpoint and exhaustion-associated genes, including PDCD1, TOX, LAG3, HAVCR2, TIGIT, CTLA4, ENTPD1, CXCL13, LAYN, and EOMES. Progenitor-associated genes, including TCF7, IL7R, CCR7, SELL, LEF1, and BACH2, were used to support interpretation of the non-effector/progenitor-exhaustion compartment. These marker programs were used for anchor selection and module-score support, but they were not used as the direct feature space for the classifier. Instead, the effector-to-non-effector axis was learned in C2S embedding space. A regularized ridge logistic model was trained on C2S embedding-derived features from effector and non-effector anchor cells, producing a continuous axis in which low scores corresponded to effector-like cells, high scores to non-effector/progenitor-exhaustion-like cells, and intermediate scores to candidate latent effector-like cells.

The primary downstream state annotation used the relaxed PCA50_Ridge model, in which C2S embeddings were reduced to the top 50 C2S principal components before ridge logistic modeling. To assess robustness, we compared this primary model with PCA100_Ridge and Full2048_ridge models. The full-dimensional (2048 C2S dimensions) model preserved the complete C2S embedding representation, whereas PCA-reduced models provided denoised lower-dimensional feature spaces. For each model, cells were assigned to operational C2S-defined states, and latent-state calls were compared across models. This analysis distinguished broadly reproducible latent/intermediate cells from model-sensitive calls and tested whether the Latent_effector compartment reflected a stable structure in C2S space rather than a feature-space-specific artifact. Cells were summarized into four operational analysis states: Effector, Latent_effector/intermediate, NonEffector/progenitor-exhaustion, and Other/ambiguous. The continuous C2S axis was treated as the primary quantitative output, while categorical labels were used for visualization and downstream summaries. In the relaxed PCA50_Ridge definition, Effector cells were assigned using an effector-side axis cutoff of ≤0.30, NonEffector cells using a non-effector-side cutoff of ≥0.70, and candidate Latent_effector cells using an intermediate axis interval of 0.30–0.70 together with effector/exhaustion module-score support. A stricter PCA50_Ridge definition used more conservative thresholds, with Effector ≤0.25, Latent_effector 0.35–0.65, and NonEffector ≥0.75. Thus, relaxed labels defined a broader candidate intermediate compartment, whereas strict labels provided a conservative sensitivity definition.

### Leave-One-patient-Out analysis

Patient-level robustness was evaluated using leave-one-patient-out analysis in the DS1 and DS2 discovery cohorts. In each iteration, one patient was held out, the ridge model was refit using the remaining discovery patients, and cells from the held-out patient were projected onto the refit axis. Full-model and leave-one-out projections were compared using axis-score correlation, state-label agreement, Latent_effector fraction, and latent-cell Jaccard overlap. This analysis tested whether the PCA50_Ridge-derived axis and state assignments were driven by any single discovery patient.

### External validation

Additional dataset (DS3) was processed independently through the same Cell Ranger, Seurat QC, sentence-generation, and C2S-embedding workflow. New cells were then projected onto the frozen DS1/DS2-trained C2S axis without redefining anchors, PCA loadings, ridge coefficients, or state thresholds. This enabled independent evaluation of latent effector-like state reproducibility, response-associated axis shifts, TCR clonotype connectivity, and pathway support.

### Visualization methods

Biological interpretation used several independent evidence layers. PHATE^24^ and UMAP were used to visualize the C2S manifold, while diffusion-based ordering was used as trajectory-like visualization rather than formal lineage inference. TCR analyses assessed paired clonotype continuity, clone expansion, state-spanning clonotypes. BEAM-T antigen-associated clones, where available, were interpreted using clone size and timepoint reproducibility. PROGENy pathway activity was inferred from the normalized gene-expression matrix^25^. Cell-level pathway scores were summarized at a pseudobulk-like level by dataset, patient, sample/timepoint, response, and C2S-defined state to reduce the effect of unequal cell numbers across samples. These summaries were used to compare signaling activity across *Effector, Latent_effector, NonEffector,* and *Other* states and to determine whether the C2S-defined Latent_effector compartment showed pathway features distinct from canonical effector and non-effector/progenitor-exhaustion programs.

---

**Suppl. Fig. 1.**
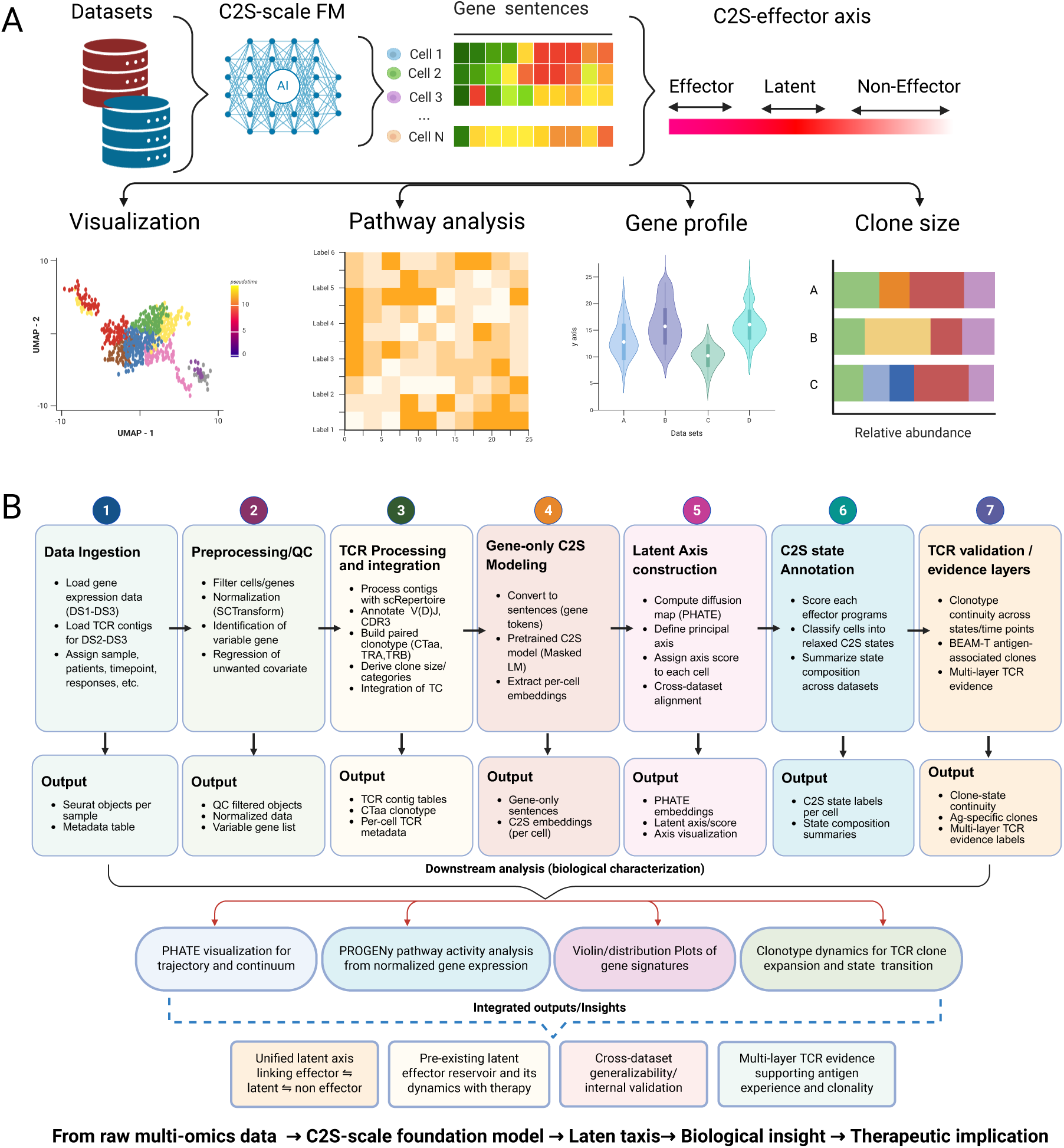
Schematic of overall strategy of this study (A) and detailed working plan of C2S-scale model architecture and analysis pipeline (B).

**Suppl. Fig. 2.**
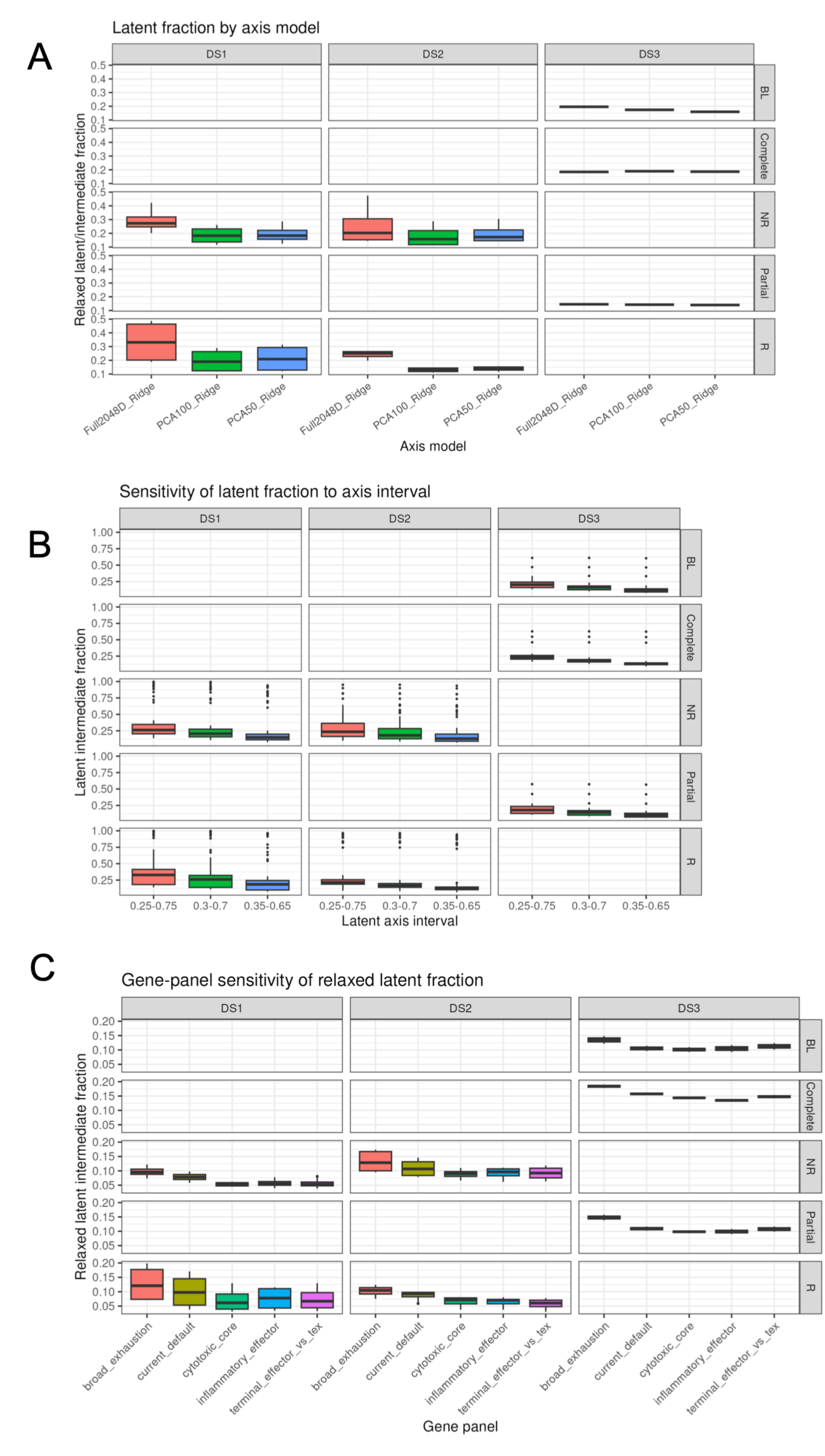
Axis-model sensitivity and latent stability. Sensitivity of latent fraction to (A) models, (B) axis interval, and (C) gene programs.

**Suppl. Fig. 3.**
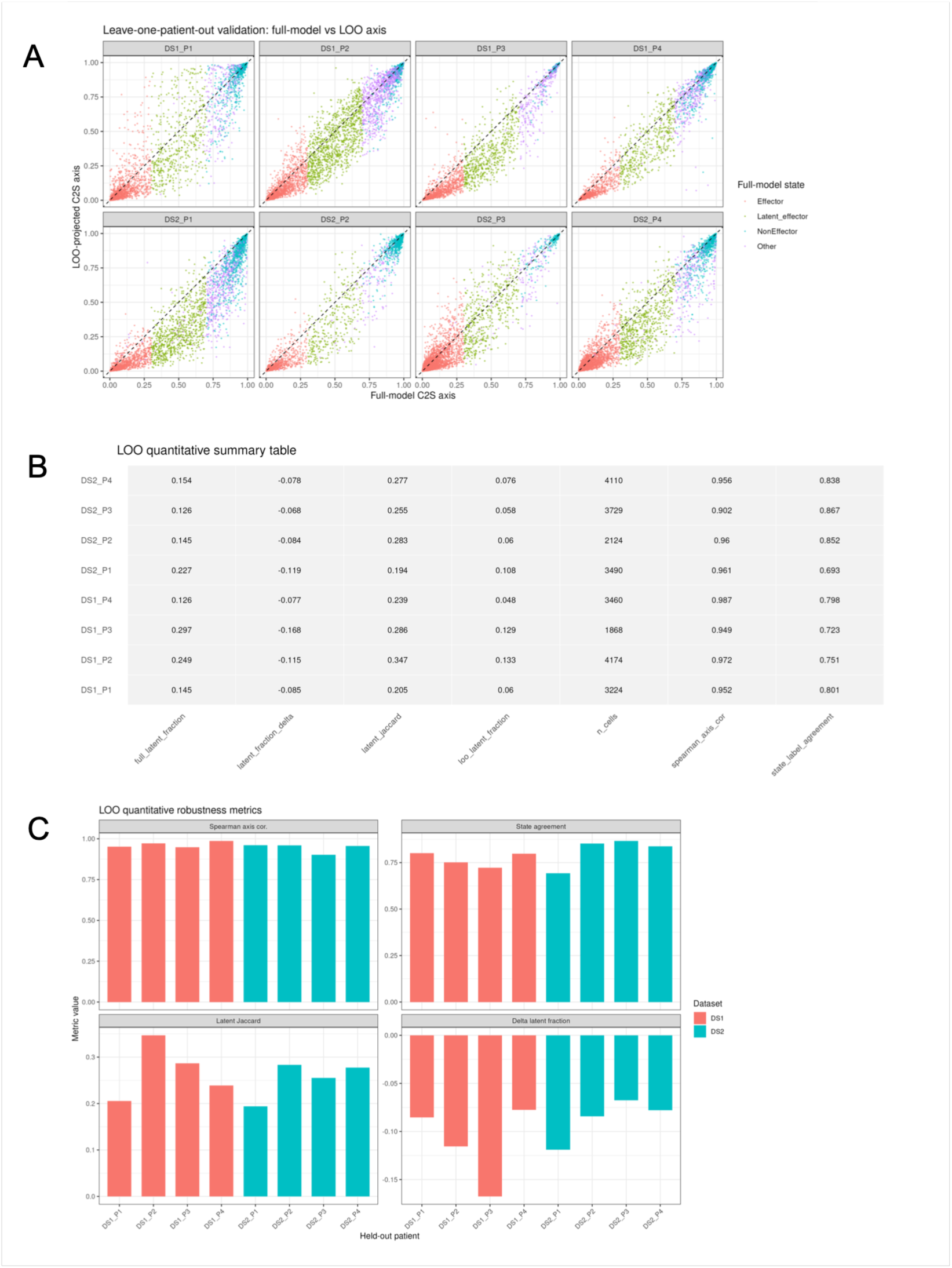
Leave-one-patient-out (LOO) robustness of the C2S axis. For each DS1 or DS2 patient, the C2S axis model was refit after excluding that patient, and held-out cells were projected onto the refit axis (A). The LOO axis preserved the expected effector-to-non-effector gradient and showed high agreement with the full-model axis. (B-C) LOO models assigned lower Latent_effector fractions than the full model, indicating that threshold-defined latent labels are more conservative under patient holdout, whereas the continuous C2S axis remains robust.

**Suppl. Figure 4.**
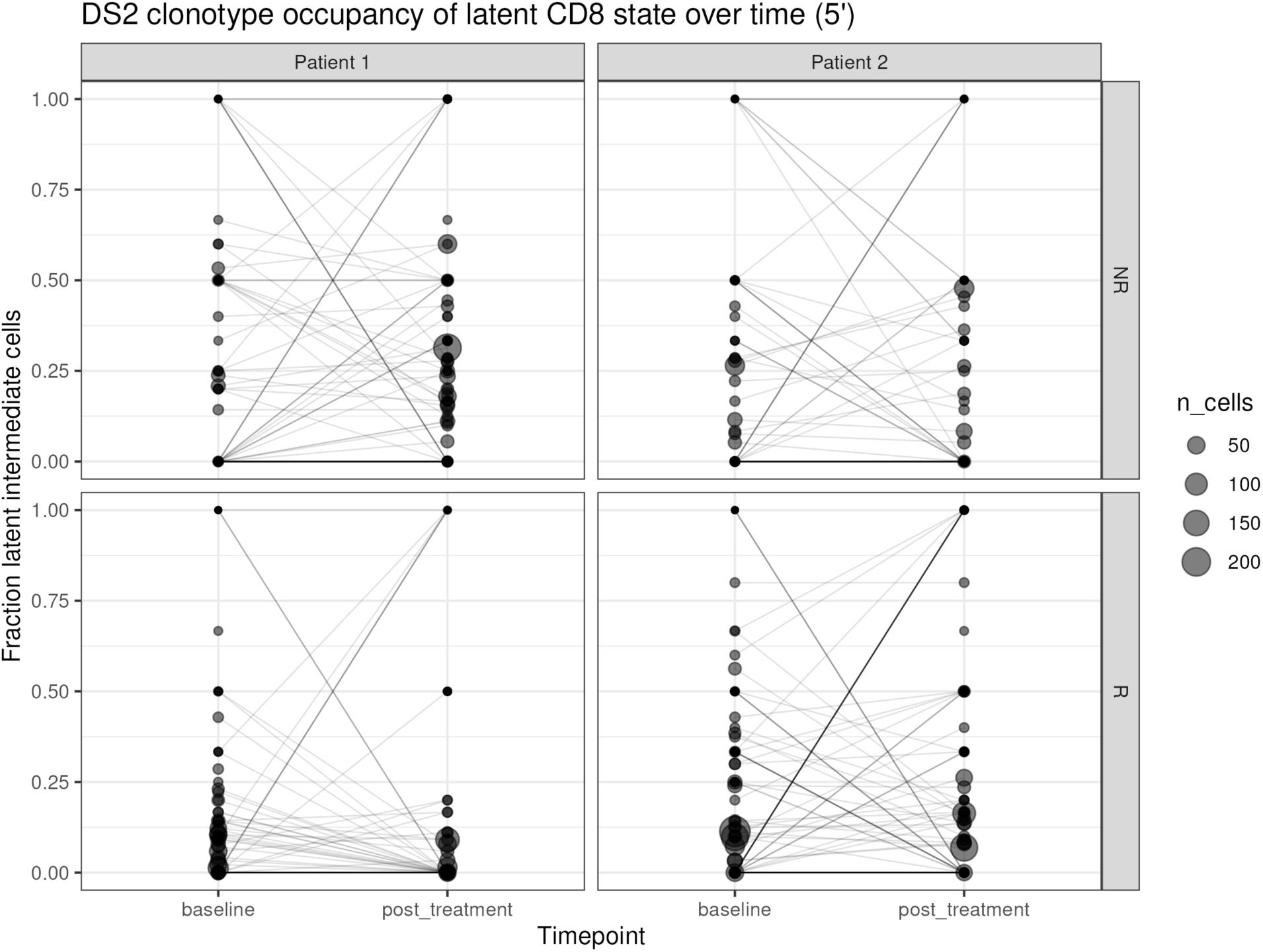
TCR clonotype occupancy of latent state of CD8^+^ T cells over time in response to anti-PD-1 therapy. The TCR clonality in fraction of latent effector state was determined by scTCR/scRNA-seq in DS2 from baseline through post therapy time points. Each line represents each TCR clones and the size of circle represents their clonal size.

**Suppl. Table S1.** The current default marker program was used for primary state annotation.

| Program# | Effector genes | Exhaustion / NonEffector genes | Progenitor genes | Use in analysis | Biological rationale / source basis |
| --- | --- | --- | --- | --- | --- |
| Current default | GZMB, PRF1, GNLY, NKG7, IFNG, CX3CR1, FGFBP2, KLRG1, CCL5, GZMA | PDCD1, TOX, LAG3, HAVCR2, TIGIT, CTLA4, ENTPD1, CXCL13, LAYN, EOMES | TCF7, IL7R, CCR7, SELL, LEF1, BACH2 | Primary analysis | Balanced cytotoxic effector, checkpoint/exhaustion, and progenitor-memory program; used for the main PCA50_Ridge state labels |
| cytotoxic_core | NKG7, GNLY, PRF1, GZMB, GZMA, GZMH, CCL5 | PDCD1, TOX, LAG3, HAVCR2, TIGIT, CTLA4, ENTPD1 | TCF7, IL7R, CCR7, SELL, LEF1 | Sensitivity analysis | Tests whether latent-state detection is driven by canonical cytotoxic machinery |
| terminal_effector_vs_tex | CX3CR1, FGFBP2, KLRG1, PRF1, GZMB, GZMH, GNLY, NKG7 | TOX, TOX2, PDCD1, HAVCR2, LAG3, TIGIT, CXCL13, LAYN, ENTPD1 | TCF7, BACH2, LEF1, IL7R, CCR7, SELL | Sensitivity analysis | Tests a sharper contrast between terminal cytotoxic effector cells and tumor-exhausted T cells |
| Broad exhaustion | GZMB, PRF1, GNLY, NKG7, IFNG, CX3CR1, FGFBP2, KLRG1, CCL5, GZMA, GZMH | PDCD1, TOX, TOX2, LAG3, HAVCR2, TIGIT, CTLA4, ENTPD1, CXCL13, LAYN, BATF, NR4A1, NR4A2, EOMES | TCF7, IL7R, CCR7, SELL, LEF1, BACH2, CD27, CD28 | Sensitivity analysis | Tests whether latent calls remain robust when exhaustion/non-effector identity is defined broadly |
| inflammatory_effector | IFNG, TNF, FASLG, CCL3, CCL4, CCL5, GZMB, PRF1, NKG7 | PDCD1, TOX, LAG3, HAVCR2, TIGIT, CTLA4, ENTPD1 | TCF7, IL7R, CCR7, SELL, LEF1, BACH2 | Sensitivity analysis | Tests whether latent-state detection is supported by inflammatory/cytokine effector activity rather than cytotoxic granule genes alone |

## Notes

### Competing Interest Statement

The authors have declared no competing interest.

## References

1. Cui H et al. Towards multimodal foundation models in molecular cell biology. Nature. 2025;640(8059):623–633. 10.1038/s41586-025-08710-y

2. Bunne C et al. How to build the virtual cell with artificial intelligence: Priorities and opportunities. Cell. 2024;187(25):7045–7063. 10.1016/j.cell.2024.11.015

3. Karr JR et al. A whole-cell computational model predicts phenotype from genotype. Cell. 2012;150(2):389–401. 10.1016/j.cell.2012.05.044

4. Rao VM et al. Generalist biological artificial intelligence in modeling the language of life. Nat Biotechnol. 2026;44(6):918–933. 10.1038/s41587-026-03064-w

5. Moor M et al. Foundation models for generalist medical artificial intelligence. Nature. 2023;616(7956):259–265. 10.1038/s41586-023-05881-4

6. Dibaeinia P et al. Virtual cells need context, not just scale. bioRxiv. 202610.64898/2026.02.04.703804

7. Rizvi SA et al. Scaling large language models for next-generation single-cell analysis. bioRxiv. 202610.1101/2025.04.14.648850

8. Philip M, Schietinger A. Cd8 t cell differentiation and dysfunction in cancer. Nature Reviews Immunology. 2022;22(4):209–223. 10.1038/s41577-021-00574-3

9. Thommen DS, Schumacher TN. T cell dysfunction in cancer. Cancer Cell. 2018;33(4):547–562. 10.1016/j.ccell.2018.03.012

10. Sade-Feldman M et al. Defining t cell states associated with response to checkpoint immunotherapy in melanoma. Cell. 2018;175(4):998–1013 e1020. 10.1016/j.cell.2018.10.038

11. van der Leun AM, Thommen DS, Schumacher TN. Cd8(+) t cell states in human cancer: Insights from single-cell analysis. Nat Rev Cancer. 2020;20(4):218–232. 10.1038/s41568-019-0235-4

12. Oliveira G et al. Phenotype, specificity and avidity of antitumour cd8(+) t cells in melanoma. Nature. 2021;596(7870):119–125. 10.1038/s41586-021-03704-y

13. Wen T et al. Nkg7 is a t-cell-intrinsic therapeutic target for improving antitumor cytotoxicity and cancer immunotherapy. Cancer Immunol Res. 2022;10(2):162–181. 10.1158/2326-6066.CIR-21-0539

14. Yan Y et al. Cx3cr1 identifies pd-1 therapy-responsive cd8+ t cells that withstand chemotherapy during cancer chemoimmunotherapy. JCI Insight. 2018;3(8):10.1172/jci.insight.97828

15. Vera Aguilera J et al. Chemo-immunotherapy combination after pd-1 inhibitor failure improves clinical outcomes in metastatic melanoma patients. Melanoma Res. 2020;30(4):364–375. 10.1097/CMR.0000000000000669

16. Dronca RS, Leontovich AA, Nevala WK, Markovic SN. Personalized therapy for metastatic melanoma: Could timing be everything? Future Oncol. 2012;8(11):1401–1406. 10.2217/fon.12.126

17. Liew AY, Li Y, Dong H. Latent effector capacity governs reversible t cell exhaustion: A mathematical model for mechanistically predictive ai in pd-1 blockade. bioRxiv. 20262026.2004.2013.717714. 10.64898/2026.04.13.717714

18. Liew AY et al. Me1 programs latent effector capacity and grounds a mathematical model of reversible t cell exhaustion. bioRxiv. 20262026.2005.2005.722814. 10.64898/2026.05.05.722814

19. Paillon N et al. Pd-1 inhibits t cell actin remodeling at the immunological synapse independently of its signaling motifs. Sci Signal. 2023;16(813):eadh2456. 10.1126/scisignal.adh2456

20. Hor JL et al. Inhibitory pd-1 axis maintains high-avidity stem-like cd8(+) t cells. Nature. 2026;649(8095):194–204. 10.1038/s41586-025-09440-x

21. Yost KE et al. Clonal replacement of tumor-specific t cells following pd-1 blockade. Nat Med. 2019;25(8):1251–1259. 10.1038/s41591-019-0522-3

22. Yamauchi T et al. T-cell cx3cr1 expression as a dynamic blood-based biomarker of response to immune checkpoint inhibitors. Nat Commun. 2021;12(1):1402. 10.1038/s41467-021-21619-0

23. Fairfax BP et al. Peripheral cd8(+) t cell characteristics associated with durable responses to immune checkpoint blockade in patients with metastatic melanoma. Nat Med. 2020;26(2):193–199. 10.1038/s41591-019-0734-6

24. Moon KR et al. Visualizing structure and transitions in high-dimensional biological data. Nat Biotechnol. 2019;37(12):1482–1492. 10.1038/s41587-019-0336-3

25. Schubert M et al. Perturbation-response genes reveal signaling footprints in cancer gene expression. Nat Commun. 2018;9(1):20. 10.1038/s41467-017-02391-6

26. Dong H et al. A dual role for nkg7 in t-cell cytotoxicity and longevity. Cancer Immunol Res. 2025OF1–OF6. 10.1158/2326-6066.CIR-25-0384

27. Ham H et al. Lysosomal nkg7 restrains mtorc1 activity to promote cd8(+) t cell durability and tumor control. Nat Commun. 2025;16(1):1628. 10.1038/s41467-025-56931-6

28. Gerlach C et al. The chemokine receptor cx3cr1 defines three antigen-experienced cd8 t cell subsets with distinct roles in immune surveillance and homeostasis. Immunity. 2016;45(6):1270–1284. 10.1016/j.immuni.2016.10.018

29. Lan X et al. Antitumor progenitor exhausted cd8(+) t cells are sustained by tcr engagement. Nat Immunol. 2024;25(6):1046–1058. 10.1038/s41590-024-01843-8

30. Gill AL et al. Pd-1 blockade increases the self-renewal of stem-like cd8 t cells to compensate for their accelerated differentiation into effectors. Sci Immunol. 2023;8(86):eadg0539. 10.1126/sciimmunol.adg0539

31. Krishna S et al. Stem-like cd8 t cells mediate response of adoptive cell immunotherapy against human cancer. Science. 2020;370(6522):1328–1334. 10.1126/science.abb9847

32. Khan O et al. Tox transcriptionally and epigenetically programs cd8(+) t cell exhaustion. Nature. 2019;571(7764):211–218. 10.1038/s41586-019-1325-x

33. Scott AC et al. Tox is a critical regulator of tumour-specific t cell differentiation. Nature. 2019;571(7764):270–274. 10.1038/s41586-019-1324-y

34. Wu TD et al. Peripheral t cell expansion predicts tumour infiltration and clinical response. Nature. 2020;579(7798):274–278. 10.1038/s41586-020-2056-8

35. Gros A et al. Pd-1 identifies the patient-specific cd8(+) tumor-reactive repertoire infiltrating human tumors. The Journal of clinical investigation. 2014;124(5):2246–2259. 10.1172/JCI73639

36. Chen DG, Xie J, Su Y, Heath JR. T cell receptor sequences are the dominant factor contributing to the phenotype of cd8(+) t cells with specificities against immunogenic viral antigens. Cell Rep. 2023;42(11):113279. 10.1016/j.celrep.2023.113279

37. Li XY et al. Nkg7 is required for optimal antitumor t-cell immunity. Cancer Immunol Res. 2022;10(2):154–161. 10.1158/2326-6066.CIR-20-0649

38. Carnevale J et al. Rasa2 ablation in t cells boosts antigen sensitivity and long-term function. Nature. 2022;609(7925):174–182. 10.1038/s41586-022-05126-w

39. Salmond RJ, Emery J, Okkenhaug K, Zamoyska R. Mapk, phosphatidylinositol 3-kinase, and mammalian target of rapamycin pathways converge at the level of ribosomal protein s6 phosphorylation to control metabolic signaling in cd8 t cells. J Immunol. 2009;183(11):7388–7397. 10.4049/jimmunol.0902294

40. Zheng T et al. Cd160 dictates anti-pd-1 immunotherapy resistance by regulating cd8(+) t cell exhaustion in colorectal cancer. Nat Cell Biol. 2025;27(9):1555–1571. 10.1038/s41556-025-01753-3

41. Abdelfatah E et al. Predictive and prognostic implications of circulating cx3cr1(+) cd8(+) t cells in non-small cell lung cancer patients treated with chemo-immunotherapy. Cancer Res Commun. 2023;3(3):510–520. 10.1158/2767-9764.CRC-22-0383

42. Zhu M et al. Understanding suboptimal response to immune checkpoint inhibitors. Adv Biol (Weinh). 2023;7(4):e2101319. 10.1002/adbi.202101319

43. Hudson WH et al. Proliferating transitory t cells with an effector-like transcriptional signature emerge from pd-1(+) stem-like cd8(+) t cells during chronic infection. Immunity. 2019;51(6):1043–1058 e1044. 10.1016/j.immuni.2019.11.002

44. Zander R et al. Cd4(+) t cell help is required for the formation of a cytolytic cd8(+) t cell subset that protects against chronic infection and cancer. Immunity. 2019;51(6):1028–1042 e1024. 10.1016/j.immuni.2019.10.009

45. Haslam A, Olivier T, Prasad V. How many people in the us are eligible for and respond to checkpoint inhibitors: An empirical analysis. Int J Cancer. 2025;156(12):2352–2359. 10.1002/ijc.35347

46. Gicobi JK, Dellacecca ER, Dong H. Resilient t-cell responses in patients with advanced cancers. Int J Hematol. 202210.1007/s12185-022-03424-7

47. Rodrigues Pessoa R et al. Circulating antigen-primed cytotoxic t-cells in patients with renal tumors treated with surgery. Urol Oncol. 2023;41(9):393 e391–393 e397. 10.1016/j.urolonc.2023.05.009

48. Miller BC et al. Subsets of exhausted cd8(+) t cells differentially mediate tumor control and respond to checkpoint blockade. Nat Immunol. 2019;20(3):326–336. 10.1038/s41590-019-0312-6

49. Daniel B et al. Divergent clonal differentiation trajectories of t cell exhaustion. Nat Immunol. 2022;23(11):1614–1627. 10.1038/s41590-022-01337-5

50. Baessler A, Vignali DAA. T cell exhaustion. Annu Rev Immunol. 2024;42(1):179–206. 10.1146/annurev-immunol-090222-110914

51. Belk JA et al. Genome-wide crispr screens of t cell exhaustion identify chromatin remodeling factors that limit t cell persistence. Cancer Cell. 2022;40(7):768–786 e767. 10.1016/j.ccell.2022.06.001

52. McLane LM, Abdel-Hakeem MS, Wherry EJ. Cd8 t cell exhaustion during chronic viral infection and cancer. Annu Rev Immunol. 2019;37(457–495. 10.1146/annurev-immunol-041015-055318

53. Zwijnenburg AJ et al. Graded expression of the chemokine receptor cx3cr1 marks differentiation states of human and murine t cells and enables cross-species interpretation. Immunity. 2023;56(8):1955–1974 e1910. 10.1016/j.immuni.2023.06.025

